# Heterogeneous local dynamics revealed by classification analysis of spatially disaggregated time series data

**DOI:** 10.1101/276006

**Authors:** T. Alex Perkins, Isabel Rodriguez-Barraquer, Carrie Manore, Amir S. Siraj, Guido España, Christopher M. Barker, Michael A. Johansson, Robert C. Reiner

## Abstract

Time series data provide a crucial window into infectious disease dynamics, yet their utility is often limited by the spatially aggregated form in which they are presented. When working with time series data, violating the implicit assumption of homogeneous dynamics below the scale of spatial aggregation could bias inferences about underlying processes. We tested this assumption in the context of the 2015-2016 Zika epidemic in Colombia, where time series of weekly case reports were available at national, departmental, and municipal scales. First, we performed a descriptive analysis, which showed that the timing of departmental-level epidemic peaks varied by three months and that departmental-level estimates of the time-varying reproduction number, *R*(*t*), showed patterns that were distinct from a national-level estimate. Second, we applied a classification algorithm to six features of proportional cumulative incidence curves, which showed that variability in epidemic duration, the length of the epidemic tail, and consistency with a cumulative normal density curve made the greatest contributions to distinguishing groups. Third, we applied this classification algorithm to data simulated with a stochastic transmission model, which showed that group assignments were consistent with simulated differences in the basic reproduction number, *R*_0_. This result, along with associations between spatial drivers of transmission and group assignments based on observed data, suggests that the classification algorithm is capable of detecting differences in temporal patterns that are associated with differences in underlying drivers of incidence patterns. Overall, this diversity of temporal patterns at local scales underscores the value of spatially disaggregated time series data.

## INTRODUCTION

Time series have been used for many years to make inferences about processes that shape infectious disease dynamics [1]. This long history has resulted in an appreciation for a number of common challenges for time series analysis [2]. One such challenge is disentangling the effects of multiple interacting forces, which can include both extrinsic forces, such as weather, and intrinsic forces, such as feedbacks associated with changes in population immunity [3,4]. An even more fundamental challenge lies in defining the time series in the first place, especially with respect to space [5]. The question is, at what spatial scale should epidemiological data be aggregated for time series analysis?

In practice, the spatial scale at which data are aggregated to form a time series is more often dictated by the scale at which data are available than by the scale that is optimal for inference or prediction. For example, during the recent invasions of chikungunya virus (CHIKV) and then Zika virus (ZIKV) across the Americas, the Pan American Health Organization published weekly case reports aggregated nationally. Despite an abundance of evidence that chikungunya and dengue viruses – which are also transmitted by *Aedes aegypti* mosquitoes – are characterized by spatially focal transmission [6,7], applications ranging from estimation of time-varying reproduction numbers [8] to forecasting [9,10] have utilized data aggregated at national scales for countries as vast and spatially heterogeneous as Brazil and Mexico.

Unlike most other countries in the Americas, routine surveillance of Zika in Colombia was reported on a weekly basis in each of its 1,123 municipalities during the 2015-2016 epidemic [11]. Although such case reports are underestimates of the true extent of transmission of most infectious diseases, particularly those with high proportions of asymptomatic infections, they still provide a uniquely valuable resource given the paucity of publicly available data at similar scales in most countries [12]. Such data are particularly valuable for Zika, given that a range of spatial scales are relevant for activities related to its prevention and control. On the one hand, vector control activities are planned and budgeted on multiple administrative levels but must be targeted on a very local level. On the other hand, communications, surveillance, and possible vaccination programs are generally planned and implemented only on larger administrative scales.

In the many contexts in which time series data are only available at larger administrative scales, analyses at those scales often assume that dynamics below that scale are homogeneous [13–15]. Our goal in this study was to test this common assumption using a unique data set on the ZIKV invasion of Colombia. We did so using a three-part approach that allowed us to characterize heterogeneity in temporal incidence patterns and to explore possible drivers of that heterogeneity. First, we performed standard descriptive analyses of epidemiological time series at three different spatial scales to characterize the extent to which results from these analyses differ across different spatial scales. Second, we performed a classification analysis of proportional cumulative incidence curves at departmental and municipal scales to identify differences in temporal dynamics at each of these scales. Third, we assessed possible drivers of these differences by validating the classification algorithm against simulated data and by exploring associations with environmental variables relevant to transmission.

As a whole, this three-part approach provides insight about heterogeneities in epidemic dynamics at different spatial scales and establishes a new method for classifying local epidemic dynamics in a manner consistent with differences in underlying drivers of those dynamics. All data and code used in this study are available at https://github.com/TAlexPerkins/TimeSeriesSpatialScale.

## METHODS

### Data

The focal point of our analysis was a collection of municipal-level time series of weekly Zika case reports at the municipal level in Colombia spanning August 2015 through September 2016. The primary source of these data was the Colombian National Institute of Health (Instituto Nacional de Salud, INS), which made official weekly reports of the cumulative numbers of suspected and confirmed Zika cases available in real time during the epidemic [16]. The version of these data that we used in this analysis were processed in a manner that addressed inconsistencies between data reported at municipal and departmental scales, as described by Siraj *et al*. [17]. Specifically, to correct for the fact that the total of municipal-level data from 2015 (3,875 cases) was less than the total of national-level data from 2015 (11,712), we imputed the 7,837 missing cases at the municipal level for 2015 by multiplying each municipality’s weekly incidence in 2015 by a factor required to achieve better known cumulative totals for each municipality as of the first week of 2016.

### Descriptive analysis of weekly case reports

We performed two preliminary analyses of differences in weekly case report patterns at different scales of spatial aggregation. First, we generated a bar plot of national case reports color-coded by which of 33 departments those national cases arose from. Likewise, for each of those departments, we generated a bar plot of departmental case reports color-coded by which of its municipalities those departmental cases arose from. Second, we made estimates of the time-varying effective reproduction number, *R*(*t*), for each time series. Following Ferguson *et al.* [8], we used the EstimateR function from the EpiEstim library [18] in R to estimate *R*(*t*) for each time series based on the method introduced by Cori *et al.* [19]. In brief, this method is based on an assumed distribution of the serial interval (i.e., the timing between onset of primary and secondary cases) that can be used to estimate the number of cases in the previous generation that gave rise to those observed in the present generation, thereby enabling estimation of *R*(*t*).

### Classification analysis of proportional cumulative incidence curves

We focused our analysis on cumulative, rather than raw, incidence because of the extreme variability in raw incidence patterns in this data set. This variability could occur for several reasons, including stochasticity [20], variability in underreporting [21], or focal transmission below the scale of spatial aggregation [22]. With raw incidence, time series with a small number of cases appear extremely noisy, and temporal patterns would be difficult to extract. With proportional cumulative incidence, vastly different temporal patterns are more readily comparable, because they all begin at 0 and end at 1 but arrive there by different paths. Others [23] have criticized the use of cumulative incidence data from epidemics, although these criticisms mostly pertain to parameter estimation and forecasting, neither of which we do here. Rather, our goal was to characterize diversity in the temporal patterns of an epidemic as viewed from different perspectives spatially.

The cumulative incidence curves that we examined were proportional, meaning that they all reached 1 at the time the last case was reported in a given area. Mathematically, for weekly reported Zika incidence *I*_*i,t*_ in location *i* in week *t*, we calculated proportional cumulative incidence as

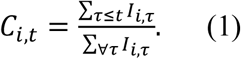

We excluded 2/33 departments and 307/1,123 municipalities from our analysis that reported no Zika cases.

As a basis for classifying proportional cumulative incidence curves, we defined six features *F* of these curves that we hypothesized represent dimensions in which curves from different areas vary. Four of these features were defined in reference to cumulative normal density curves, *Ĉ*_*i*_(*t*) that we fitted to each *C*_*i,t*_. This involved estimating mean and standard deviation parameters of *Ĉ*_*i*_ (*t*) for each *C*_*i,t*_ on the basis of least squares using the optim function in R. These six features (defined in Table 1) were chosen because they provided a way to quantify the duration of local epidemics (small *F*_*SD*_, short *F*_*Δt*_ = short epidemic), to capture whether epidemics appeared strongly locally driven (low 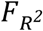, large *F*_*0*_= sporadic transmission fueled by importation), and to characterize shapes that deviated substantially from those predicted by simple epidemic models (*F*_5%_ and *F*_95%_ near zero = “SIR-like” epidemic). Although these idealized scenarios motivated the selection of these features, the fact that all six features were calculated for each *C*_*i,t*_ meant that we were able to capture a wide range of patterns in between these extremes.

**Table 1.** Features used to classify proportional cumulative incidence curves.

| Feature | Definition | Interpretation |
| --- | --- | --- |
| $F_{SD}$ | Standard deviation of $\hat{C}_i(t)$ | Rate of increase in incidence |
| $F_{R^2}$ | $R^2$ of $\hat{C}_i(t)$ in reference to $C_{i,t}$ | Degree to which incidence conforms to a smooth, epidemic-like pattern |
| $F_{5\%}$ | Difference between the 5% quantile of $C_{i,t}$ and the 5% quantile of $\hat{C}_i(t)$ | Degree to which there was a long, slow build-up in incidence |
| $F_{95\%}$ | Difference between the 95% quantile of $C_{i,t}$ and the 95% quantile of $\hat{C}_i(t)$ | Degree to which there was a long, drawn-out tail in incidence |
| $F_{\Delta t}$ | Weeks between first and last non-zero $I_{i,t}$ | Overall duration of the epidemic |
| $F_0$ | Weeks with $I_{i,t} = 0$ between first and last non-zero $I_{i,t}$ | Consistency of incidence over the course of the epidemic |

We explored variation in *C*_*i,t*_ at both departmental and municipal scales. To describe how variation in *C*_*i,t*_ curves at those scales was distributed across the six-dimensional feature space, we performed a partitioning around medoids (PAM) clustering analysis [24] on centered and scaled values of the features using the pam function in the cluster library [25] in R. This algorithm identifies medoids of *k* groups, where each medoid is a member of its group that minimizes the dissimilarity between itself and all other group members. We performed this analysis for values of *k* ranging two to ten and compared groupings for different values of *k* on the basis of their average silhouette values. A silhouette value describes how much more dissimilar one point is from points in the next most similar group compared to points in its own group [26]. An ideal classification would be indicated by silhouette values for data points in all groupings close to 1. Silhouette values nearer to or below 0 indicate that points do not cluster well with the group to which they are assigned.

### Elucidation of driving processes

To aid in the interpretation of the classification analysis of empirical patterns of temporal incidence, we performed identical analyses of simulated patterns of temporal incidence. The value of doing so is that it provides a form of validation of the classification analysis: i.e., demonstrating that it is capable of identifying groups that correspond to known differences in underlying drivers of incidence patterns. For this analysis, we defined groups of municipalities on the basis of whether the simulated *R*_0_ value for a municipality was above or below 1, given the significance of this threshold for determining invasion outcomes. We performed classification analyses on 100 data sets simulated with a stochastic model of ZIKV transmission developed by Ferguson *et al*. [8] and tailored to Colombia as described in Appendix S1. Although this model was not fitted to empirical data and is therefore limited in its realism, we did ensure that total incidence nationally was comparable to the observed national total of 85,353 suspected cases. We felt that this was important to ensure that the level of stochasticity in the simulated data was comparable to that in the empirical data. Otherwise, it was not critical that simulated data matched the empirical data, as the goal of this analysis was to evaluate the extent to which curves assigned to one group or another were classified in a way that was consistent with differences in the processes that generate those curves (i.e., *R*_0_ < 1 or > 1).

Although the analysis of simulated data provides a test of the algorithm, it does not facilitate inference of whether there truly are differences in the processes underlying local incidence patterns. Doing so convincingly would require more comprehensive analyses, ideally involving data about variables assumed to play an intermediary role in a hypothesized causal pathway between environmental variables and disease incidence [27]. To explore whether there might at least be perceptible associations between environmental variables and groups identified by the classification analysis, we performed a series of one-way analyses of variance at both departmental and municipal scales. Specifically, our objective was to examine whether mean values of relevant environmental variables differed across these groups. Variables that we examined included modeled values of *R*_0_ derived from Perkins *et al*. [28] as described in Appendix S1, and seven variables compiled for municipalities and departments in Colombia by Siraj *et al*. [17]: *Ae. aegypti* occurrence probability, two measures of normalized difference vegetation index (NDVI), mean temperature, percent urban land cover, human population, and the gross cell product (GCP), a spatially disaggregated version of the gross domestic product economic index.

## RESULTS

### Descriptive analysis of weekly case reports

As a whole, the temporal pattern at the national level was consistent with what could be construed as a typical epidemic trajectory, marked by an increase over approximately five months, a peak around the beginning of February 2016, and a steady decline thereafter over a period of approximately eight months (Fig. 1A). Under a standard set of assumptions about epidemic dynamics, this pattern can be used to estimate the temporal trajectory of the effective reproduction number, *R*(*t*) [19]. Applying this technique at the national level yielded estimates of *R*(*t*) that began high (range: 1.5-3.5 for the first four months) and gradually declined below 1 by the time the epidemic concluded (Fig. 1A), which could be consistent with expectations for an epidemic of an immunizing pathogen in an immunologically naive host population, among other explanations.

**Figure 1.**
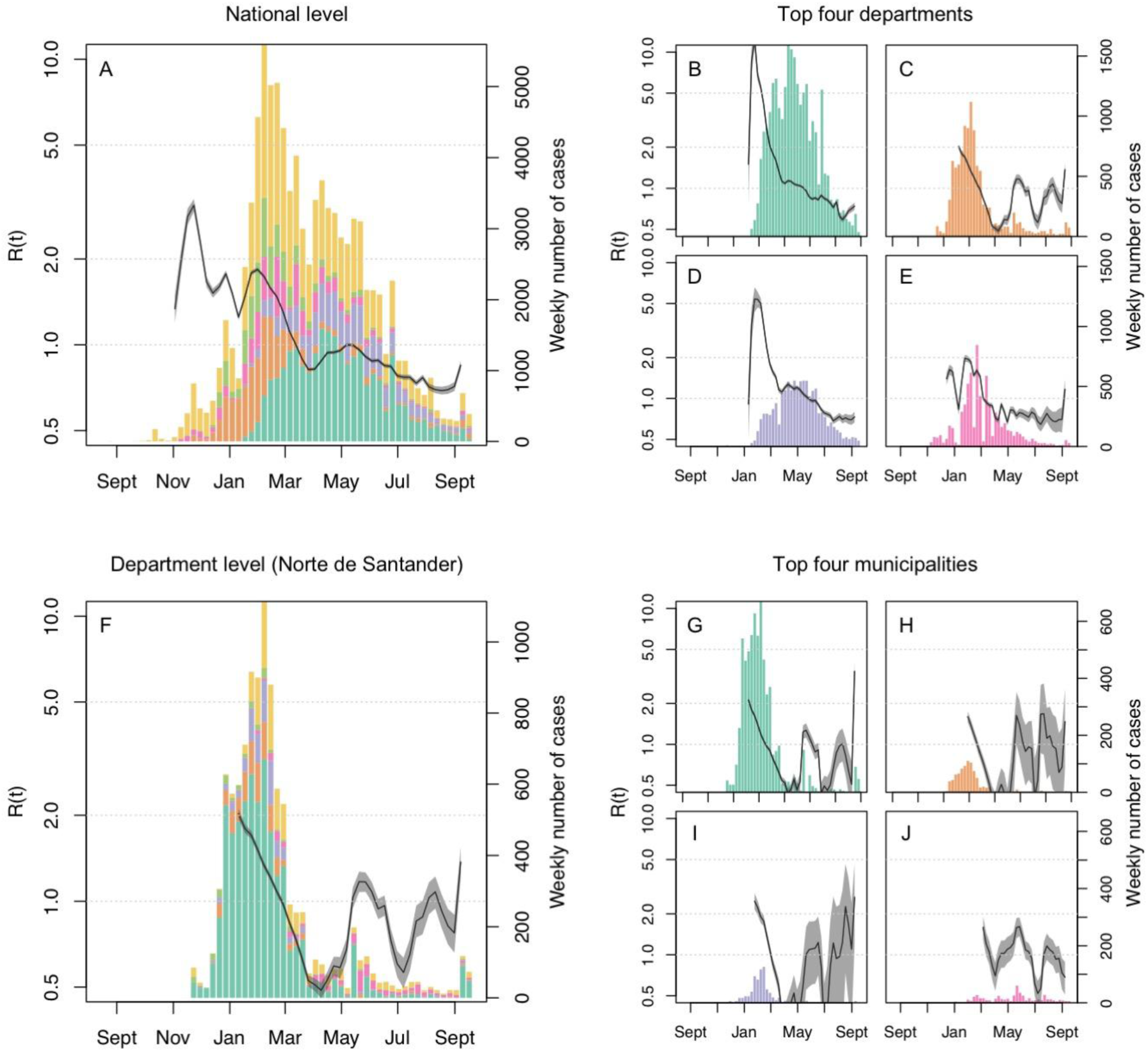
Weekly Zika case reports at the national level (A), for each of the four departments with the largest case report totals (B: Valle del Cauca; C: Norte de Santander; D: Santander; E: Tolima), at the departmental level for Norte de Santander (F), and for each of its four municipalities with the largest case report totals (G: Cucuta; H: Villa del Rosario; I: Los Patios; J: Ocaña). On the top row, colors match across A and B-E, with the addition of yellow in A that includes all departments other than those in B-E. On the bottom row, colors match across F and G-J, with the addition of yellow in F that includes all municipalities other than those in G-J. Time-varying estimates of the effective reproduction number, *R*(*t*), are shown in each panel. A map showing the location of these departments is available in Fig. S1.

Examination of temporal incidence patterns for each of the four largest departments in terms of total incidence (Valle del Cauca, Norte de Santander, Santander, Tolima, Fig. S1) showed that patterns at the departmental level were quite different than those at the national level. First, the timing of peak incidence in the departments in Fig. 1B-1E varied by around three months. Second, the shapes of the incidence patterns in those departments varied, with Valle del Cauca and Santander (Fig. 1B & 1D) showing high incidence sustained over a period of several months and Norte de Santander and Tolima (Fig. 1C & 1E) showing sharper peaks trailed by relatively low incidence for several months after.

This high degree of variability in temporal incidence patterns had substantial impacts on estimates of *R*(*t*). At the national level, *R*(*t*) estimates never exceeded 3.5, whereas in Santander *R*(*t*) was estimated to exceed 5 (Fig. 1D) and in Valle del Cauca it was estimated to exceed 10 (Fig. 1B), due in both cases to more rapid increases in incidence at the departmental level than the national level. In Norte de Santander, *R*(*t*) appeared to twice fall well below 1 but then quickly rise back above 1 (Fig. 1C).

Examination of temporal patterns at the municipal scale revealed even more variability in temporal patterns than at the department level. In the department of Norte de Santander (Fig. 1C), for example, it was clear that one municipality dominated the departmental pattern (Fig. 1F). The municipalities with the second and third highest incidence both experienced short, unimodal patterns of incidence during the first two months, but incidence patterns thereafter were mostly low and erratic (Fig. 1G & 1H). Other municipalities in the department had only low, erratic incidence with no sign of a distinct epidemic (e.g., Fig. 1J). With the exception of the first few weeks of transmission, estimates of *R*(*t*) at the municipal level were characterized by erratic fluctuations and much more uncertainty than was apparent at the departmental or national level.

### Classification analysis of proportional cumulative incidence curves

At the departmental level, there was only modest clustering overall, with the highest average silhouette value corresponding to two groups (0.256), a slightly lower value for three groups (0.254), and falling no lower than 0.201 for up to ten groups (Fig. S2). Example fits of cumulative normal density curves for clustering based on two groups are available in Fig. S3. *F*_*SD*_ and *F*_95%_ were the features that were most important for distinguishing two groups (Fig. S4), and *F*_*Δt*_ contributed further to distinguishing three groups (Fig. S5). Differences in *F*_*SD*_ were associated with a difference of approximately two months in the time elapsed between the attainment of 5% and 80% of cumulative incidence (Fig. 2, top left: blue longer than red), and differences in *F*_95%_ were associated with a difference of approximately two months in the time elapsed between the attainment of 80% and 99% of cumulative incidence, but for different groups (Fig. 2, top left: red longer than blue). Overall, this meant that the time elapsed between attainment of 5% and 99% of cumulative incidence for both groups was similar, but with one group experiencing epidemics that were fast initially but slow to finish and another group experiencing epidemics that were slower initially but finished more quickly. These patterns were clearest for the curves associated with the medoid of each group (Fig. 2, top) but were generally apparent for the curves associated with the groups as a whole (Fig. S6). Spatially, groups tended to cluster along northern, central, and southern strata (Fig. 3, left), with incidence-weighted cartographs showing that the epidemic was mostly dominated by distinct northern and central strata (Fig. 3, top right).

**Figure 2.**
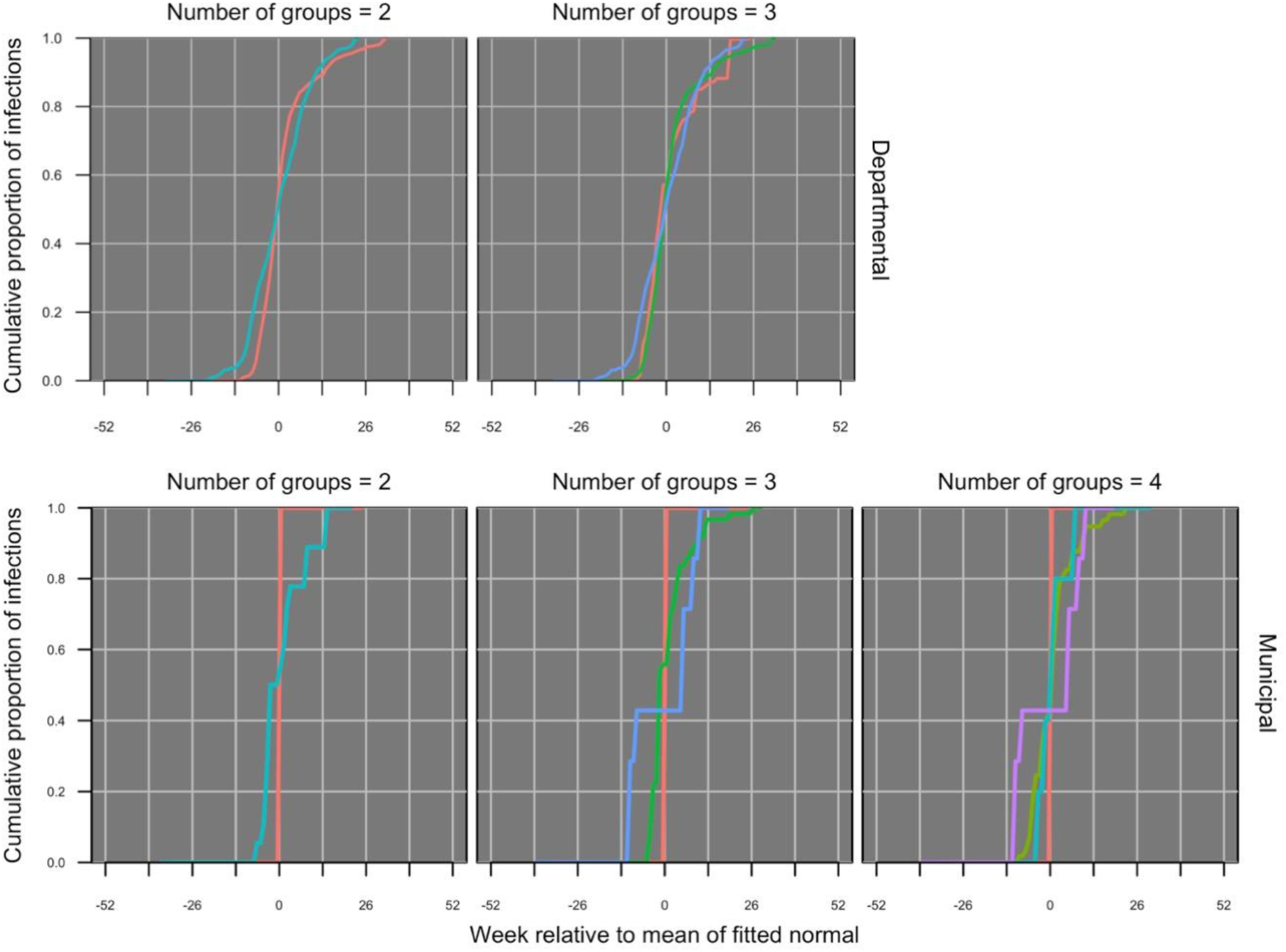
Proportional cumulative incidence curves at the departmental level (top) with two (left) or three (right) groups and at the municipal level (bottom) with two (left), three (middle), and four (right) groups. Only one representative curve is shown for each group, with that curve being chosen on the basis of being associated with the medoid of its group.

**Figure 3.**
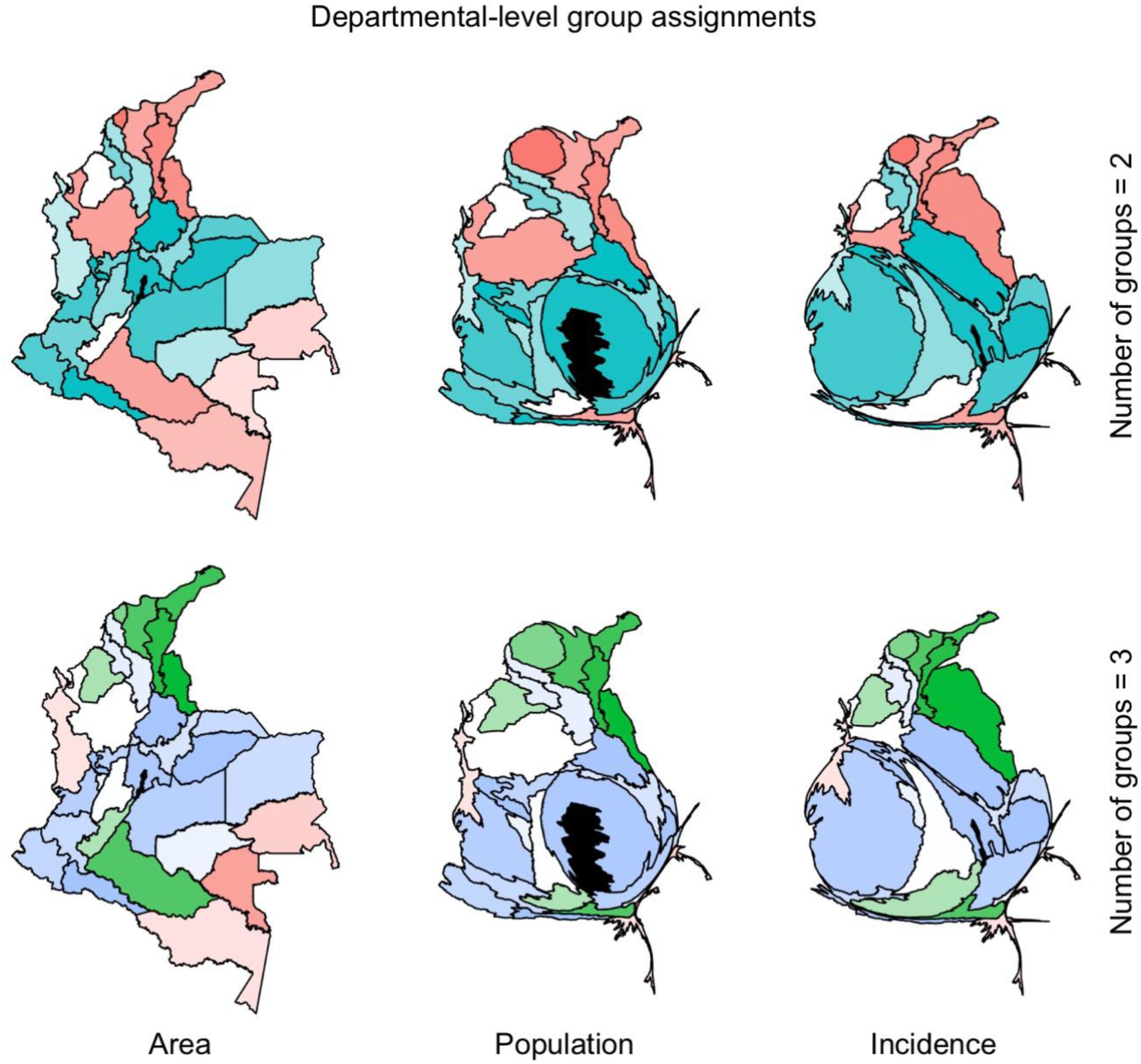
Cartograms at the departmental level weighted by area (left), population (center), and incidence (right). Department assignments to two (top) and three (bottom) groups are indicated by color, with transparency inversely proportional to silhouette value. The one department (Bogotá) with zero incidence is indicated in black and given a weight equivalent to 1/5 of a case to allow for its inclusion in the right column.

There was somewhat stronger clustering at the municipal level, with the highest average silhouette value corresponding to three groups (0.352), somewhat lower values for five and six groups (0.334, 0.326), and no lower than 0.297 for up to ten groups (Fig. S7). Example fits of cumulative normal density curves for clustering based on three groups are available in Fig. S8. *F*_*Δt*_ and *F*_*SD*_ were the features that were most important in distinguishing two groups (Fig. S9), *F*_95%_ made additional contributions to distinguishing three groups (Fig. S10), and 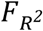 contributed to distinguishing four groups (Fig. S11). Proportional cumulative incidence curves with short *F*_*Δt*_ and small *F*_*SD*_ comprised the most visually distinct group and remained relatively consistent regardless of the number of groups (Fig. 2, bottom). Some differences among the other groups were also apparent in the proportional cumulative incidence curves, with some having a long tail (Fig. 2, bottom middle: green) or two discrete jumps (Fig. 2, bottom middle: blue). The timing of discrete jumps varied across municipalities, but curves within a group otherwise resembled the curve associated with the medoid for that group (compare Fig. 2 bottom with Fig. S12). Spatially, departments generally consisted of a mixture of municipalities from different groups, and the prominence of some groups in the cartograms varied depending on whether the cartograms were weighted by area, population, or incidence (Fig. 4). The cartograms weighted by population showed that a sizeable portion of the population lives in cities that had no reported cases, such as Medellín and Bogotá (Fig. 4, black in the center column). Among municipalities that did have reported cases, the cartograms weighted by incidence showed that a relatively large proportion of reported cases came from municipal-level epidemics characterized by large *F*_*Δt*_ and *F*_*SD*_ (Fig. 4, right column).

**Figure 4.**
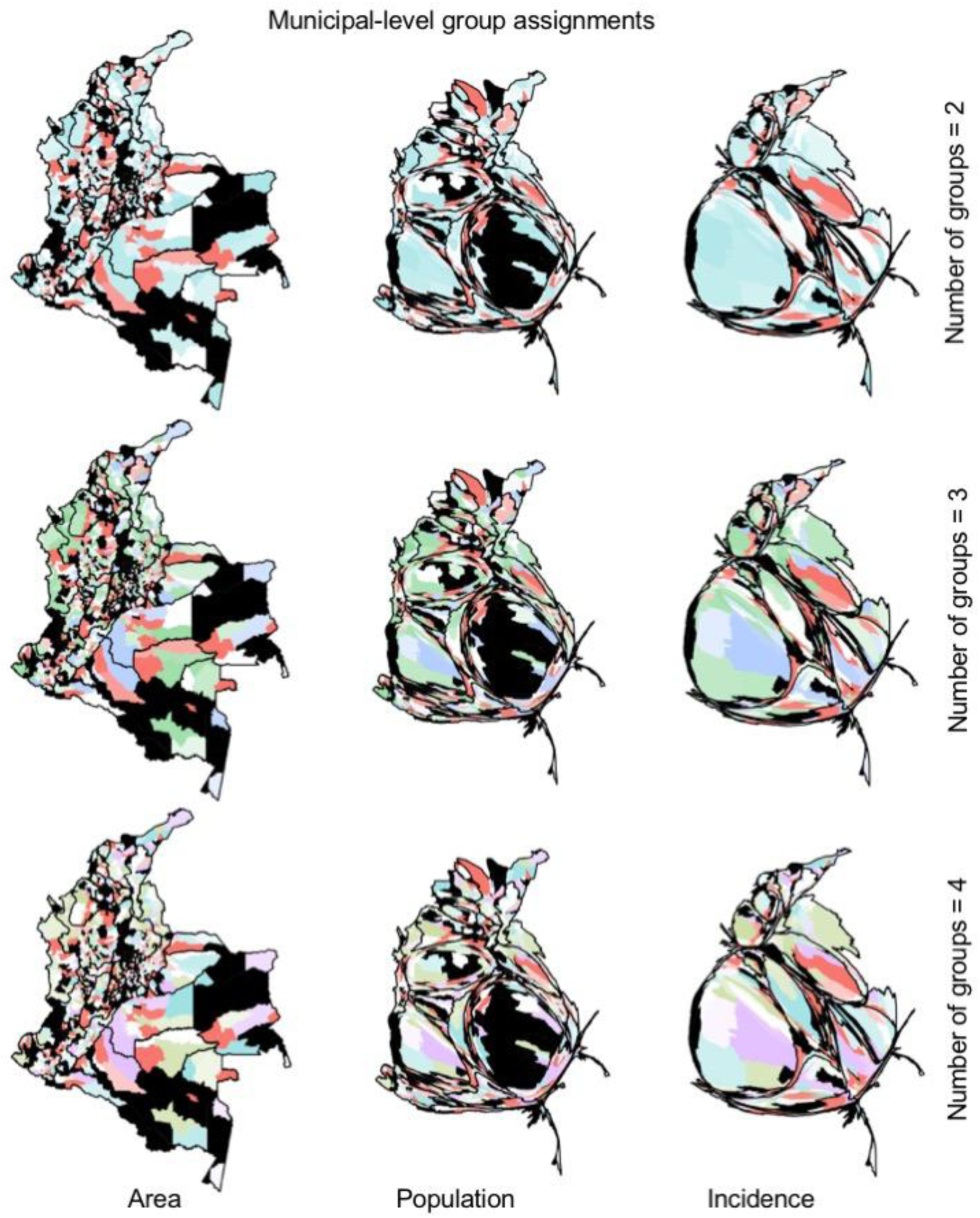
Cartograms at the municipal level weighted by area (left), population (center), and incidence (right). Municipality assignments to two (top), three (middle), and four (bottom) groups are indicated by color, with transparency inversely proportional to silhouette value.

Municipalities with zero incidence are indicated in black and were given a weight equivalent to 1/5 of a case to allow for their inclusion in the right column.

### Elucidation of driving processes

We focused our analysis of simulated data at the municipal level given that the simulation model was not equipped to simulate transmission between municipalities, which is likely important for recreating departmental-level patterns. Overall, our model parameterization assumed that *R*_0_ > 1 in 34.6% of municipalities. A total of 12.6% (range: 10.4-14.1%) of municipalities had zero simulated cases, with 99.0% (range: 97.0-100.0%) of those having *R*_0_ < 1.

Out of 100 simulated datasets, the classification algorithm selected two groups eight times, three groups 80 times, and five and six groups four times each. Average silhouette value was 0.313 (range: 0.288-0.347) when there were two groups and 0.327 (range: 0.291-0.352) when there were three groups (see Fig. S13 for a representative silhouette plot from a randomly selected simulated dataset). Although this indicates a modest preference of the algorithm for three groups, we focused subsequent analyses on the two-group classification due to our desire to evaluate the correspondence between groups selected by the classification analysis and groups defined by *R*_0_ above or below 1.

With the two-group classification, 99.1% (range: 90.3-100.0%) of municipalities with *R*_0_ > 1 were placed into the group characterized by larger *F*_*Δt*_ and *F*_*SD*_. Of the municipalities with *R*_0_ < 1, 74.0% (range: 36.3-80.5%) were also placed into that group, with the others placed into the group with smaller *F*_*Δt*_ and *F*_*SD*_ (see Fig. S14 for an example from a randomly selected simulated dataset). When municipalities were classified into three groups, a new group characterized by moderately low *F*_*Δt*_ and *F*_*SD*_ and negative *F*_95%_ contained 18.8% (range: 0.2-36.1%) of municipalities with *R*_0_ > 1 and 44.7% (range: 23.0-56.5%) with *R*_0_ < 1 (see Fig. S15 for an example from a randomly selected simulated dataset). In the presence of this third group, 79.9% (range: 63.4-89.7%) of municipalities with *R*_0_ > 1 and 32.1% (range: 22.8-38.8%) with *R*_0_ < 1 were placed into the group characterized by larger *F*_*Δt*_ and *F*_*SD*_

Visual inspection of five simulated datasets showed that the proportional cumulative incidence curves of municipalities placed in the group characterized by large *F*_*Δt*_ and *F*_*SD*_ generally resembled the curves of municipalities with *R*_0_ > 1 (Fig. 5, red). In contrast, proportional cumulative incidence curves of municipalities with *R*_0_ < 1 were more diverse than those placed in the group characterized by low *F*_*Δt*_ and *F*_*SD*_ (Fig. 5, blue). A similar pattern was apparent spatially, with municipalities placed in the group characterized by large *F*_*Δt*_ and *F*_*SD*_ generally overlapping with municipalities with *R*_0_ > 1, but municipalities with *R*_0_ < 1 frequently placed in the group characterized by large *F*_*Δt*_ and *F*_*SD*_ (Fig. 6).

**Figure 5.**
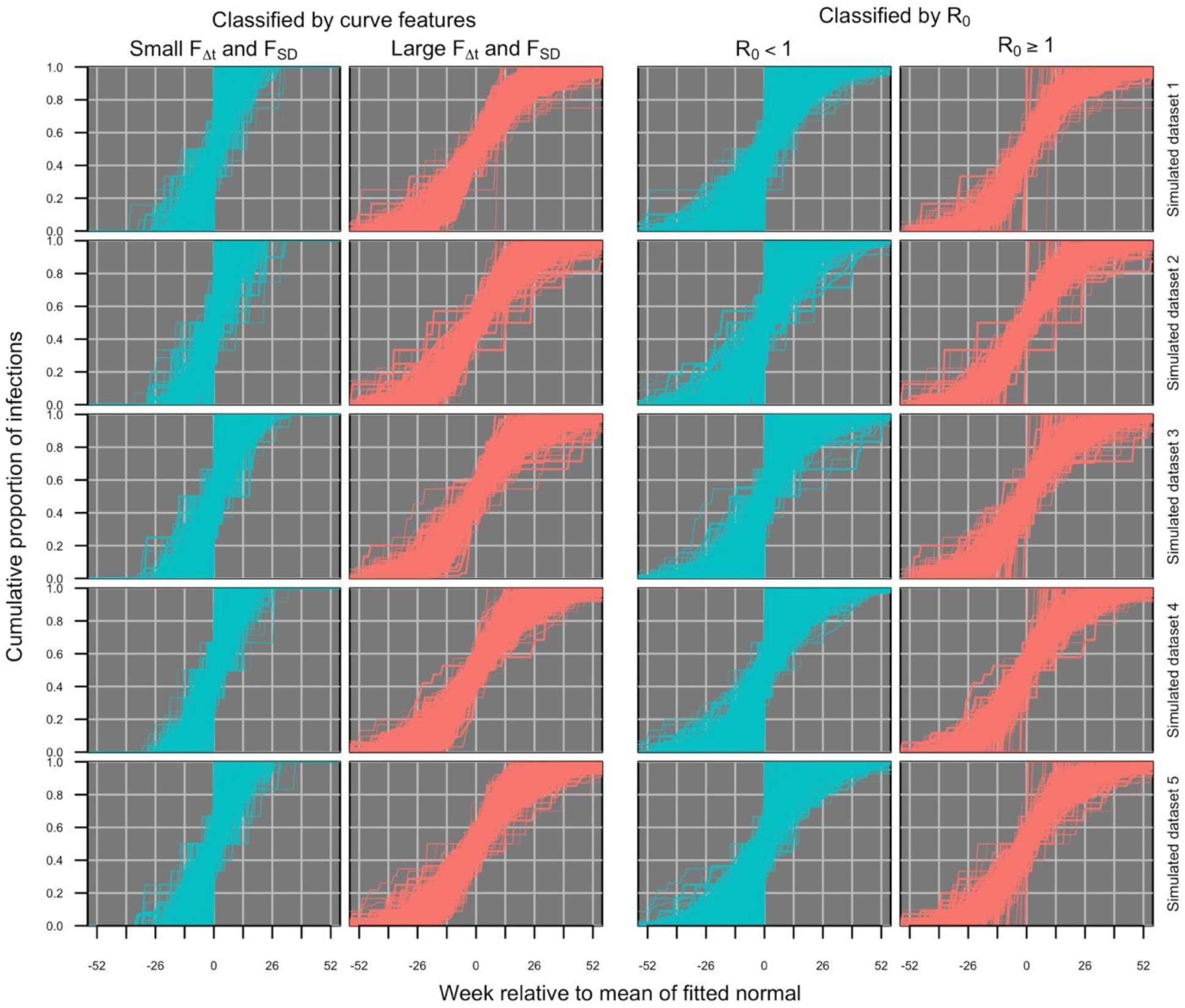
Proportional cumulative incidence curves at the municipal level from five randomly selected simulated datasets. The left two columns show two different groups classified by the curve classification algorithm, and the right two columns show two different groups defined by whether those municipalities have a *R*_0_ above or below 1.

**Figure 6.**
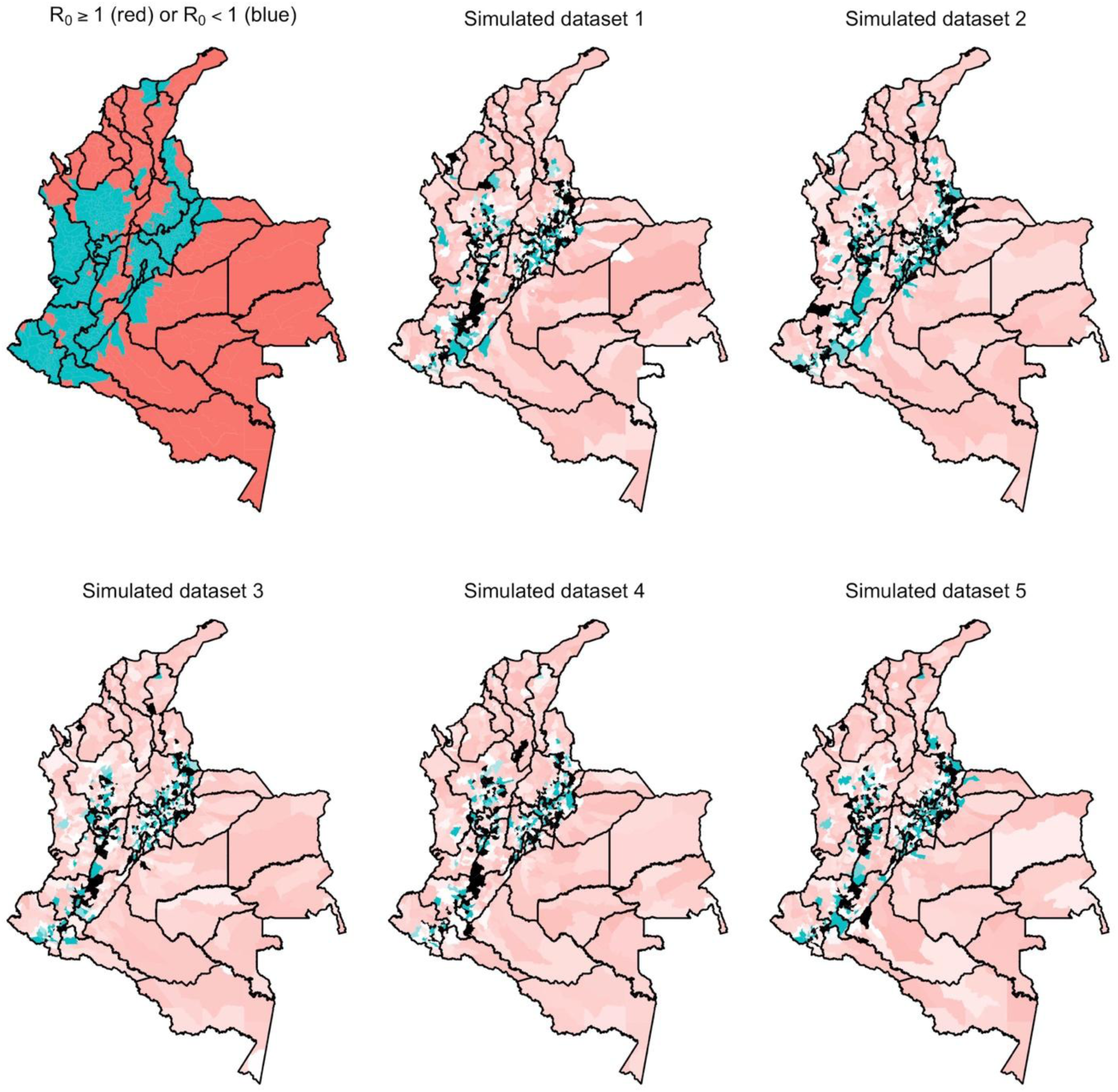
Cartograms at the municipal level weighted by area based on five randomly selected simulated datasets. Each municipality’s status as having *R*_0_ > 1 (red) or *R*_0_ < 1 (blue) is indicated in the top left panel. In each of five simulated datasets shown in the other panels, municipality assignments to two groups are indicated by color, with transparency inversely proportional to silhouette value.

Such a high proportion of municipalities with *R*_0_ < 1 being placed into the group characterized by large *F*_*Δt*_ and *F*_*SD*_ may be an artefact of imported case patterns being forced unrealistically strongly according to national-level incidence patterns (Appendix S1). This gave populous municipalities with low *R*_0_ the appearance of an epidemic more characteristic of a municipality with higher *R*_0_. Consequently, the classification algorithm may perform better on empirical data from municipalities with *R*_0_ < 1 than this analysis of simulated data suggests.

With respect to the empirical data, group assignments at the municipal scale were associated with perceptible differences in relevant environmental variables. For two groups, differences between groups were statistically significant for all eight variables examined (*p*<0.002 for all; Table S1). The group typified by steep, short curves (Fig. 2, bottom left: red) was associated with lower *Ae. aegypti* occurrence probability (0.04 vs. 0.05; *F*_836_=19.7, *p*<10^-5^), higher NDVI (aqua: 0.09 vs. 0.07; *F*_836_=12.9, *p*<10^-3^) (terra: 0.10 vs. 0.07; *F*_836_=13.3, *p*<10^-3^), lower temperature (21.7 vs. 23.9 °C; *F*_836_=32.9, *p*<10^-7^), lower urban cover (0.02 vs. 0.07; *F*_836_=29.6, *p*<10^-7^), lower population (13,506 vs 49,144; *F*_836_=10.2, *p*<10^-2^), lower GCP (6,016 vs. 6,676; *F*_836_=8.3, *p*<10^-2^), and lower *R*_0_ (1.1 vs. 1.7; *F*_836_=16.1, *p*<10^-4^) (Table S1). Differences among groups were significant only for the urban cover variable for three groups, and for no variables for four groups (Table S1). At the departmental scale, group assignments based on empirical data were generally not associated with differences in relevant environmental variables (Table S2).

## DISCUSSION

Temporal incidence patterns play a vital role in modeling infectious disease dynamics and inferring drivers thereof. By analyzing data from the 2015-2016 Zika epidemic in Colombia, we showed that temporal patterns can appear very different depending on the spatial scale at which data are aggregated. Whereas national-level dynamics appeared to follow a unimodal pattern consistent with behavior of standard epidemic models, departmental-level dynamics were somewhat more varied and municipal-level dynamics were the most varied. Simulations of our transmission model suggest that high variability in municipal-level dynamics results from differences in *R*_0_ > 1 or < 1, as well as the stochasticity of transmission dynamics in populations of this size. Combining our observations of empirical patterns at different spatial scales with a formal classification of temporal incidence patterns and a model-based exploration of mechanisms capable of generating those patterns, we deduced that there is distinct variation in temporal patterns subnationally and that much of that variation may be driven by spatial variation in local conditions. Associations between group assignments and relevant environmental variables were most apparent at the municipal scale, consistent with the hypothesis that linkages between temporal dynamics and underlying processes are strongest at fine spatial scales.

Similar to our findings of differing dynamics at municipal and departmental scales, theoretical analyses of a range of ecological models have proposed that dynamics approach deterministic behavior as spatial scales grow larger and data become increasingly more aggregated [29]. For example, methods based on long-term dynamics have been proposed for identifying the scales at which behavior transitions from stochastic to deterministic in models of plant competition and predator-prey interactions [30,31]. Epidemics, however, are inherently transient in nature, leaving open the question of how best to define characteristic spatial scales in that context. It is certainly the case that the data from Colombia that we examined displayed greater stochasticity at finer spatial scales. At the same time, the greater variability in temporal patterns that we observed at finer scales suggests that models that aspire to a deterministic representation of behavior at coarser scales must account for spatial structure at finer scales. Indeed, a recent attempt to fit a national-scale transmission model to national-scale time series of Zika case reports from Colombia showed that ignoring subnational spatial structure inhibited that model’s fit to the data [32]. A theoretical exploration of similar issues concluded that the scale at which spatial structure must be modeled explicitly is expected to vary by pathogen and geographic context, with less mobile pathogens requiring explicit spatial representation at finer scales [33].

Both stochasticity and spatial interaction are expected to contribute to variability in temporal dynamics at local scales [34]. For some municipalities, temporal incidence patterns appeared to be dominated by stochasticity (e.g., those with discrete jumps). For others, there were implications for a role of spatial interaction (e.g., those with two sharp increases or a long tail). Whereas our simulation model was realistic with respect to demography and the inclusion of spatiotemporal variability in local transmission, it made the very simplistic assumption about spatial interaction that importation patterns have identical timing and magnitude in all municipalities. This may have caused municipalities with *R*_0_ < 1, particularly those with larger populations, to display patterns that simply reflected the national trend used to drive importation. Analyses of subnational spatiotemporal dynamics in a range of contexts show that importation patterns vary substantially over time and as a function of regional connectivity or being positioned on an international border [35–38]. Future work that includes more realistic spatial interaction among subnational units would be helpful for resolving the hypothesis proposed here about the importance of spatial interaction in shaping temporal patterns at each of the spatial scales that we considered.

Our analysis identified intriguing differences in temporal patterns across spatial scales, but at the same time there are important limitations to acknowledge. First, although our conclusions are not dependent on the magnitude of transmission, they do require that patterns in case report data reflect patterns in underlying transmission. With a high rate of asymptomatic infection and the likelihood of extensive variability in reporting rates [39], particularly at the municipal level, some caution is due. Second, our ability to ascribe meaning to the groups identified by our classification algorithm was limited by the simplicity of our simulation model, particularly with respect to spatial interaction. Consequently, while this analysis identified important relationships between spatial scale and epidemic characteristics, it does not provide a complete or comprehensive understanding of the spatial transmission dynamics of ZIKV in Colombia. Third, our model relied on a simplified description of seasonal transmission, when in fact patterns of seasonality could vary spatially and interact with introduction timing [40].

Previous analyses of Zika [8,32], as well as chikungunya [9,41], have drawn inferences and made forecasts on the basis of nationally aggregated time series data. These efforts depend on the implicit assumption that spatially disaggregated temporal patterns are homogeneous and consistent with spatially aggregated temporal patterns. Our analysis showed that while national-level patterns may be somewhat reflective of departmental-level patterns, municipal-level patterns of cumulative incidence are diverse and not well approximated by national-level patterns. This finding presents a challenge, given that data at such a fine scale are often not available. One potential remedy to this challenge is to specify a transmission model at a finer scale than the scale at which data are available and then aggregate modeled incidence patterns prior to fitting to aggregated data. Taking this approach to modeling the chikungunya epidemic in Colombia demonstrated that a model specified at a higher spatial resolution than the scale at which data were available provided a better fit to aggregated data than a model specified at the scale at which data were aggregated [42]. Similar approaches will likely be necessary to understand spatial variation in transmission dynamics for Zika, which remains important for time-sensitive applications such as site selection for vaccine trials [43,44] and anticipating future epidemics [8].

## ACKNOWLEDGEMENTS

This research was supported by a RAPID grant from the National Science Foundation (DEB 1641130) and by a Young Faculty Award from the Defense Advanced Research Projects Agency (D16AP00114).

## Appendix S1. Description of simulation model of Zika virus transmission in Colombia

We simulated data sets comparable to the observed data using an R implementation of the ZIKV transmission model described by Ferguson *et al.* [8] parameterized to match the municipal-level *R*_0_ values described by Perkins *et al*. [28], which are informed by municipal-level temperature, mosquito occurrence probability, and GCP. The model by Ferguson *et al.* [8] had a number of attractive features, including plausible values of a number of parameters common to ZIKV transmission models, realistic accounting of the timing of transmission-relevant processes in mosquitoes and humans, seasonal variation in transmission, and the ability to capture multiple forms of stochasticity associated with transmission and surveillance. In brief, the model assumes that humans transition from a susceptible compartment into a recovered and immune compartment following a period of incubation and infectiousness and that mosquitoes become infectious and remain so following bites of infectious humans and a seasonally variable incubation period. Mosquito population density is also seasonally variable, driven by seasonal variation in larval carrying capacity and adult mortality. A full description of the model can be found in the paper by Ferguson *et al.* [8].

To drive the model, we based estimates of the basic reproduction number, *R*_0_, on a set of ZIKV epidemic size projections for Latin America made early in the epidemic using relationships between environmental variables and transmission metrics [28]. To obtain a single value of *R*_0_ for each municipality, we took a weighted sum of the *R*_0_ raster at 5 km x 5 km resolution weighted by a raster layer of human population projections [45] aggregated to that scale by Siraj *et al*. [17]. We calibrated these *R*_0_ estimates to observed dynamics in Colombia by scaling municipal values of *R*_0_ from [28] by a constant (2.72) such that the value for the municipality of Girardot, Colombia, matched an estimate of 4.61 derived from an analysis of temporal incidence patterns there [46]. The environmental variables that drove spatial variation in these *R*_0_ values include temperature, *Ae. aegypti* occurrence probability, and the gross cell product economic index.

To apply this model to Colombia, we used municipal-level human population sizes derived from WorldPop [45] and adjusted seasonally averaged mosquito densities such that seasonally averaged values of *R*_0_ matched our municipal-level *R*_0_ estimates. Another departure from the original model by Ferguson *et al*. [8] that we made was to remove explicit spatial coupling, given the complexity of doing so realistically for all 1,123 municipalities in Colombia. Instead, we simulated imported infections (i.e., infections acquired outside a given municipality) to occur at a daily per capita rate that was proportional to a normal probability density function fitted to the temporal pattern of national-scale incidence (timing of national-scale incidence: mean = 32.57 weeks after the first reported case, standard deviation = 8.85 weeks). Although this approach was not able to capture differences in the timing of importation patterns across municipalities, none of the six features of the cumulative incidence curves that we analyzed depended on the timing of the epidemic in one municipality relative to another. To approximately match the national total of 85,353 suspected Zika cases, the time-varying ZIKV importation function that we used was scaled by a value of 1.55 × 10^-3^. This value was obtained by trial-and-error tuning of example simulations in which a reporting rate of 11.5% was assumed [47]. Also, given that our interest was in short-term dynamics rather than long-term dynamics as in [8], we removed human age stratification from the model.

## SUPPORTING TABLES

**Table S1.** Summary of results from one-way analyses of variance at the municipal scale (*n*=836). For each relevant environmental variable (rows), we performed an analysis of variance to test for differences in the mean of that variable across two, three, or four groups identified by the classification analysis (columns). The *F* statistic and *p* value of each test is shown.

|  | Two groups |  | Three groups |  | Four groups |  |
| --- | --- | --- | --- | --- | --- | --- |
| | $F_{836}$ | $p$ | $F_{836}$ | $p$ | $F_{836}$ | $p$ |
| *Ae. aegypti* | 19.7 | $9.9 \times 10^{-6}$ | $8.7 \times 10^{-3}$ | 0.93 | 0.86 | 0.35 |
| NDVI<sub>terra</sub> | 13.3 | $2.8 \times 10^{-4}$ | 1.1 | 0.30 | 0.046 | 0.83 |
| NDVI<sub>aqua</sub> | 12.9 | $3.5 \times 10^{-4}$ | 1.1 | 0.29 | 0.072 | 0.79 |
| Mean temp. | 32.9 | $1.4 \times 10^{-8}$ | 0.14 | 0.71 | 1.7 | 0.19 |
| Pct. urban | 29.6 | $7.0 \times 10^{-8}$ | 7.9 | $5.2 \times 10^{-3}$ | 0.029 | 0.86 |
| Population | 10.2 | $1.4 \times 10^{-3}$ | 1.7 | 0.19 | 0.50 | 0.48 |
| GCP | 8.3 | $4.1 \times 10^{-3}$ | 2.0 | 0.16 | 1.6 | 0.21 |
| $R_0$ | 16.1 | $6.6 \times 10^{-5}$ | 0.056 | 0.81 | 1.7 | 0.19 |

**Table S2.**
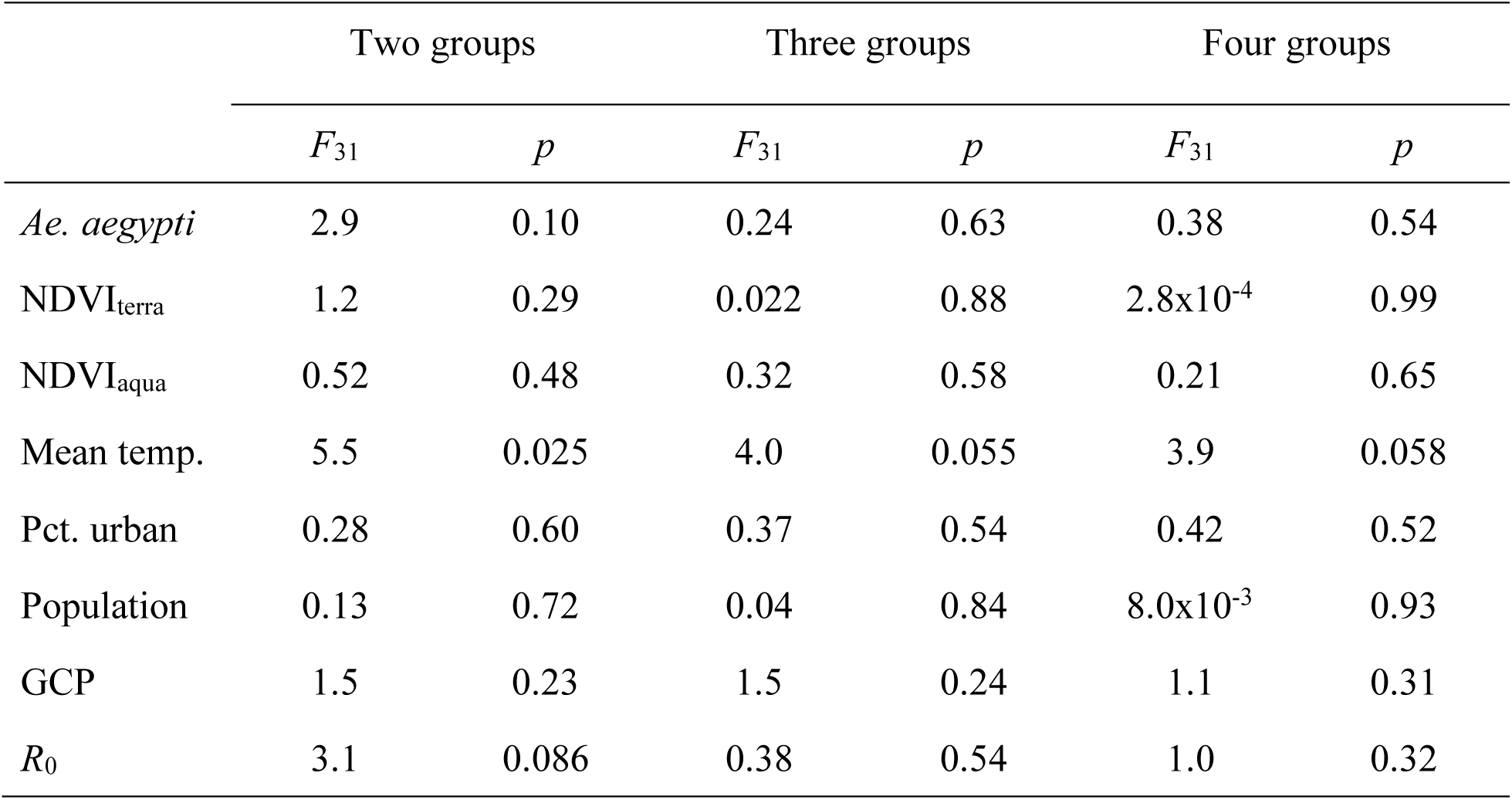
Summary of results from one-way analyses of variance at the departmental scale (*n*=31). For each relevant environmental variable (rows), we performed an analysis of variance to test for differences in the mean of that variable across two, three, or four groups identified by the classification analysis (columns). The *F* statistic and *p* value of each test is shown.

|  | Two groups |  | Three groups |  | Four groups |  |
| --- | --- | --- | --- | --- | --- | --- |
| | $F_{31}$ | $p$ | $F_{31}$ | $p$ | $F_{31}$ | $p$ |
| *Ae. aegypti* | 2.9 | 0.10 | 0.24 | 0.63 | 0.38 | 0.54 |
| NDVI<sub>terra</sub> | 1.2 | 0.29 | 0.022 | 0.88 | 2.8x10<sup>-4</sup> | 0.99 |
| NDVI<sub>aqua</sub> | 0.52 | 0.48 | 0.32 | 0.58 | 0.21 | 0.65 |
| Mean temp. | 5.5 | 0.025 | 4.0 | 0.055 | 3.9 | 0.058 |
| Pct. urban | 0.28 | 0.60 | 0.37 | 0.54 | 0.42 | 0.52 |
| Population | 0.13 | 0.72 | 0.04 | 0.84 | 8.0x10<sup>-3</sup> | 0.93 |
| GCP | 1.5 | 0.23 | 1.5 | 0.24 | 1.1 | 0.31 |
| $R_0$ | 3.1 | 0.086 | 0.38 | 0.54 | 1.0 | 0.32 |

## SUPPORTING FIGURES

**Figure S1.**
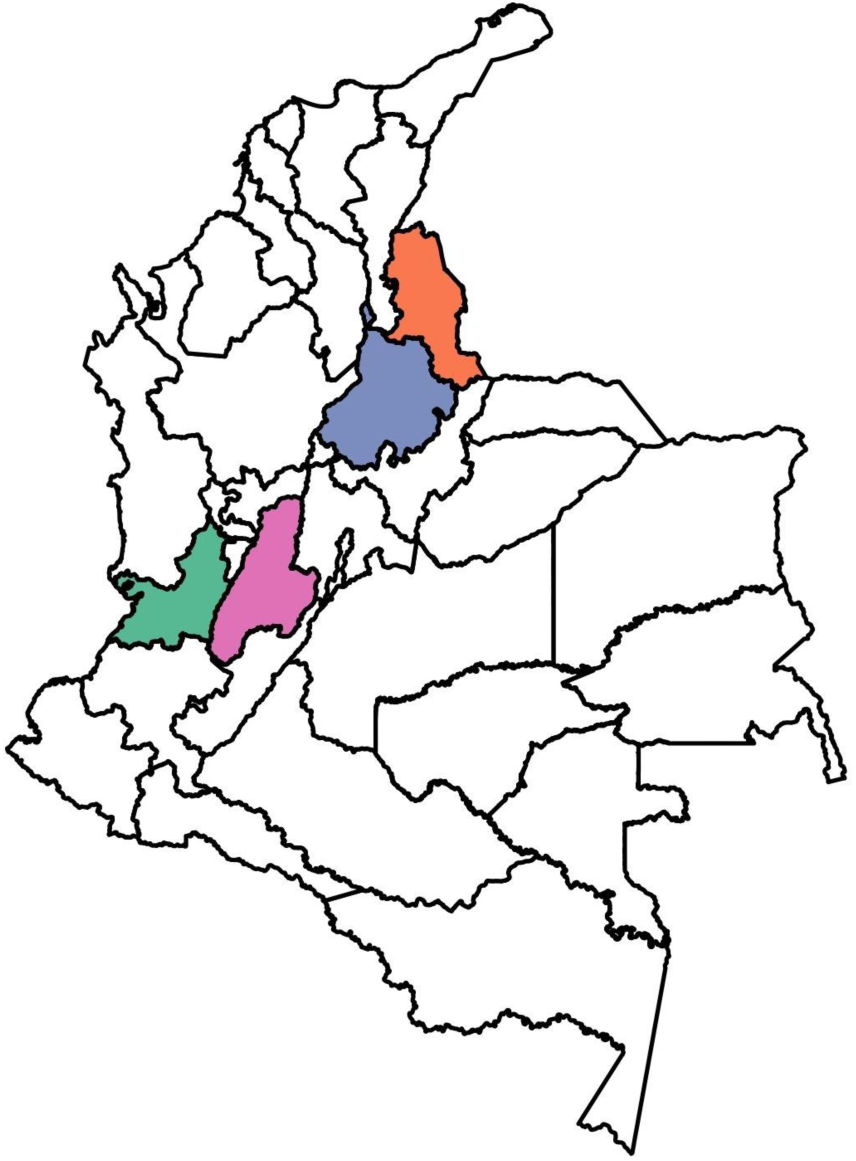
Map indicating the departments highlighted in Fig. 1 that have the four highest cumulative incidences. In order from first to fourth highest incidence, colors correspond to departments as follows: green = Valle del Cauca; orange = Norte de Santander; purple = Santander; pink = Tolima.

**Figure S2.**
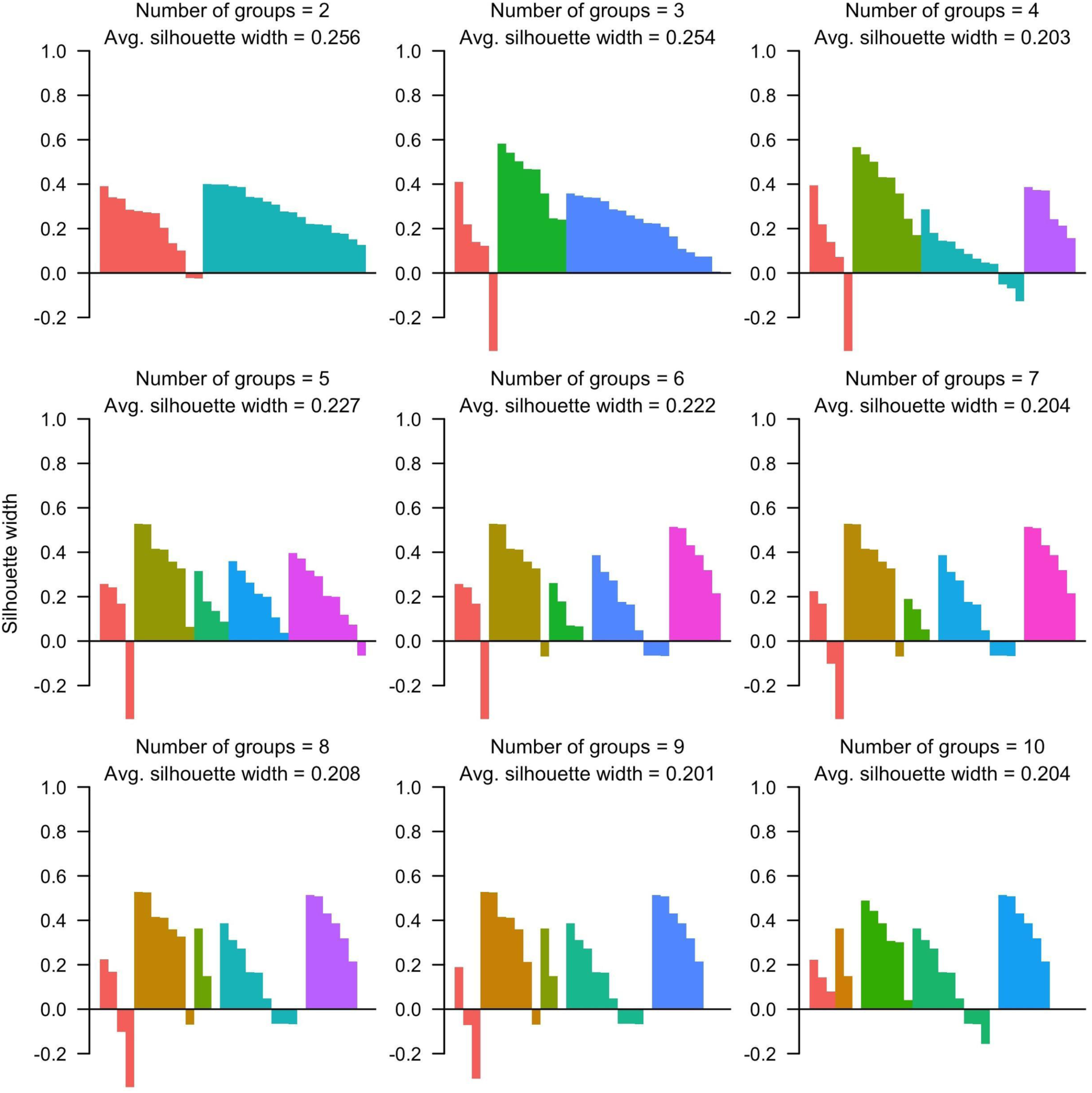
Silhouette plots at the departmental level for groups numbering two to ten obtained by partitioning around medoids. Each bar corresponds to the silhouette value of a given department according to the group assignments indicated by different colors in each panel. Higher average silhouette values indicate stronger clustering.

**Figure S3.**
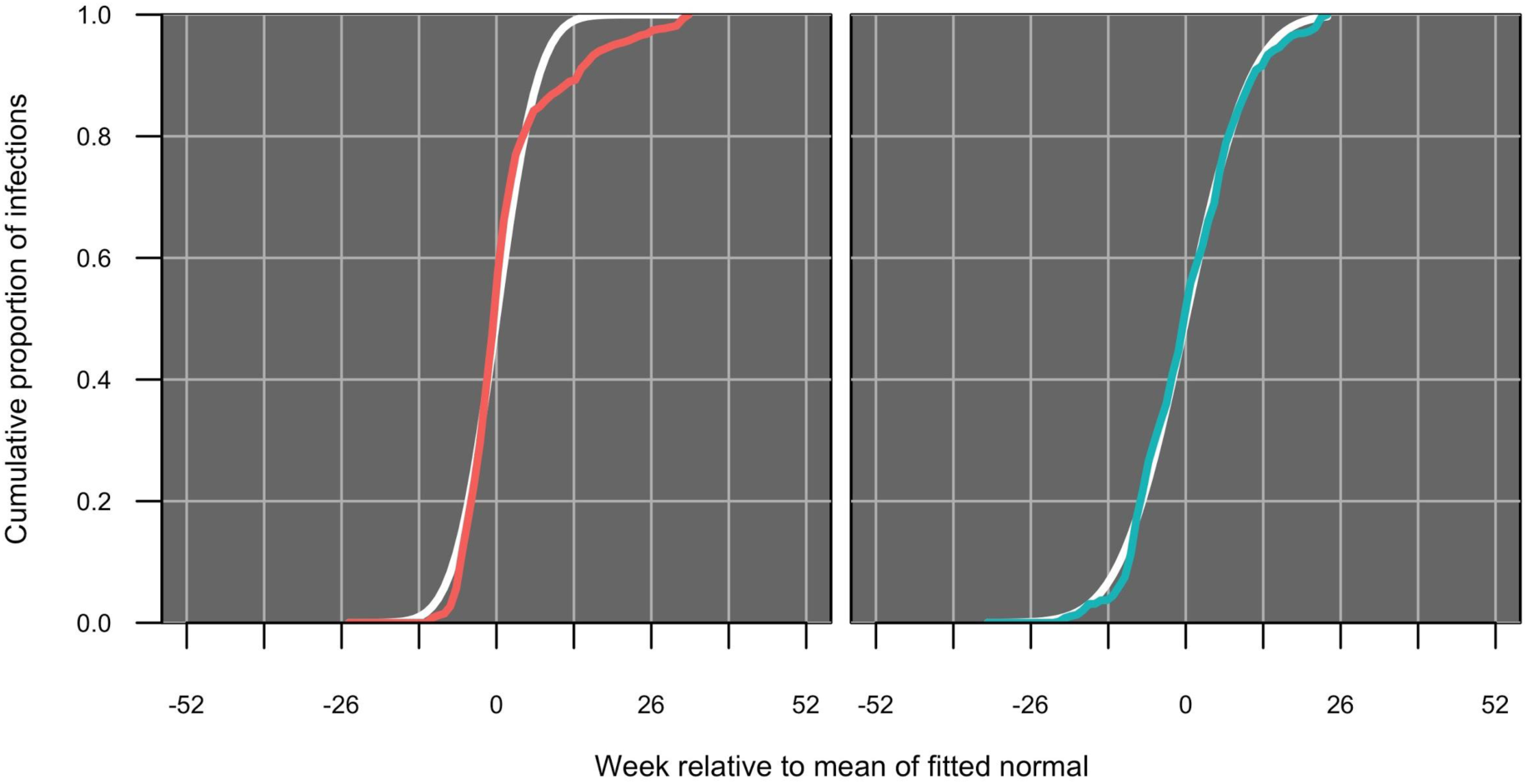
Proportional cumulative incidence curves (red, blue) shown against cumulative normal density curves (white) for medoids of the classification based on two groups at the departmental level.

**Figure S4.**
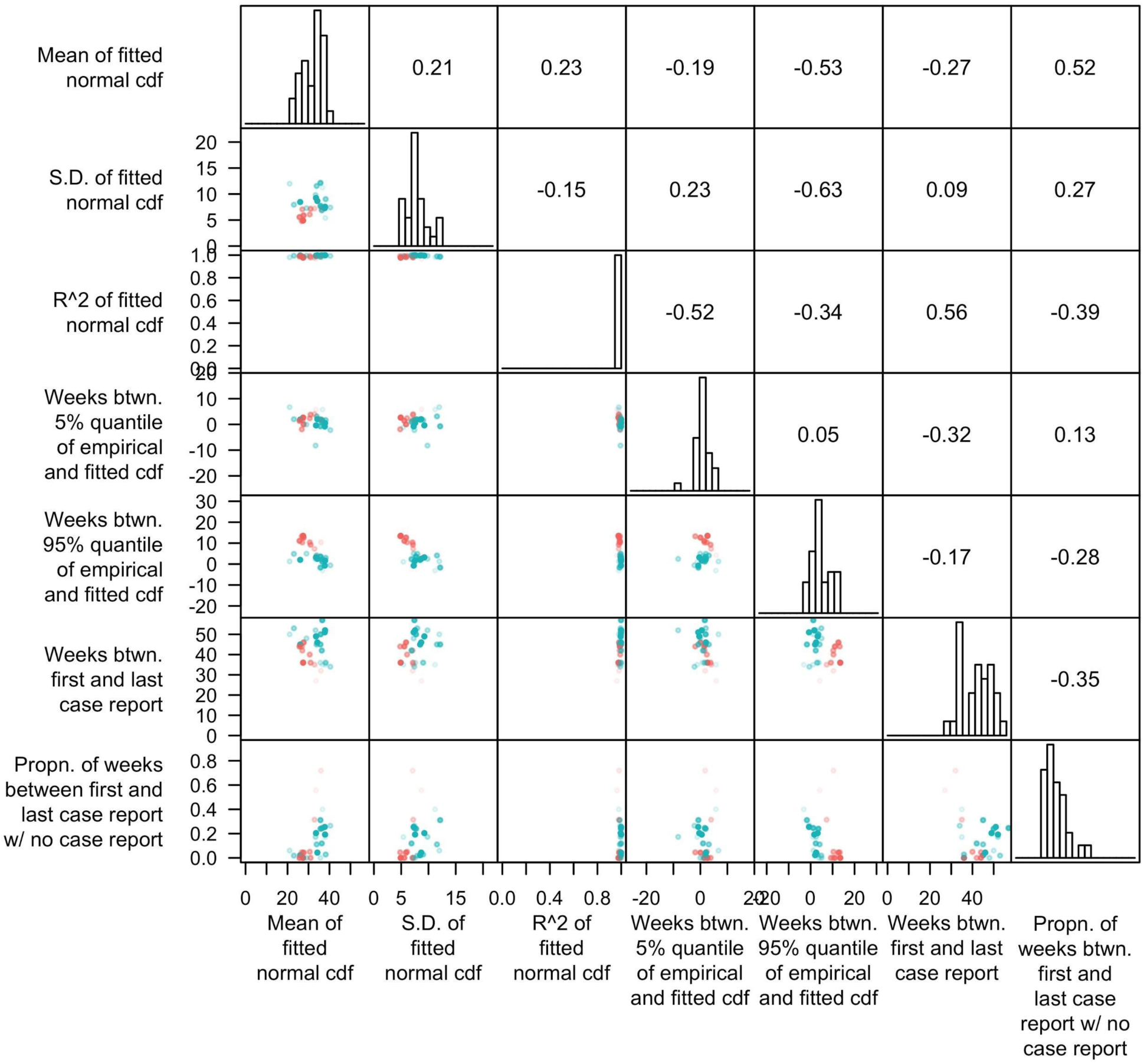
Pairwise plots of features of proportional cumulative incidence curves, with colors distinguishing group assignment of the departments into one of two groups. Histograms show the marginal distributions of the features, and numbers in the upper right half indicate pairwise correlation coefficients between each pair of features. The transparency of each point is inversely proportional to silhouette value.

**Figure S5.**
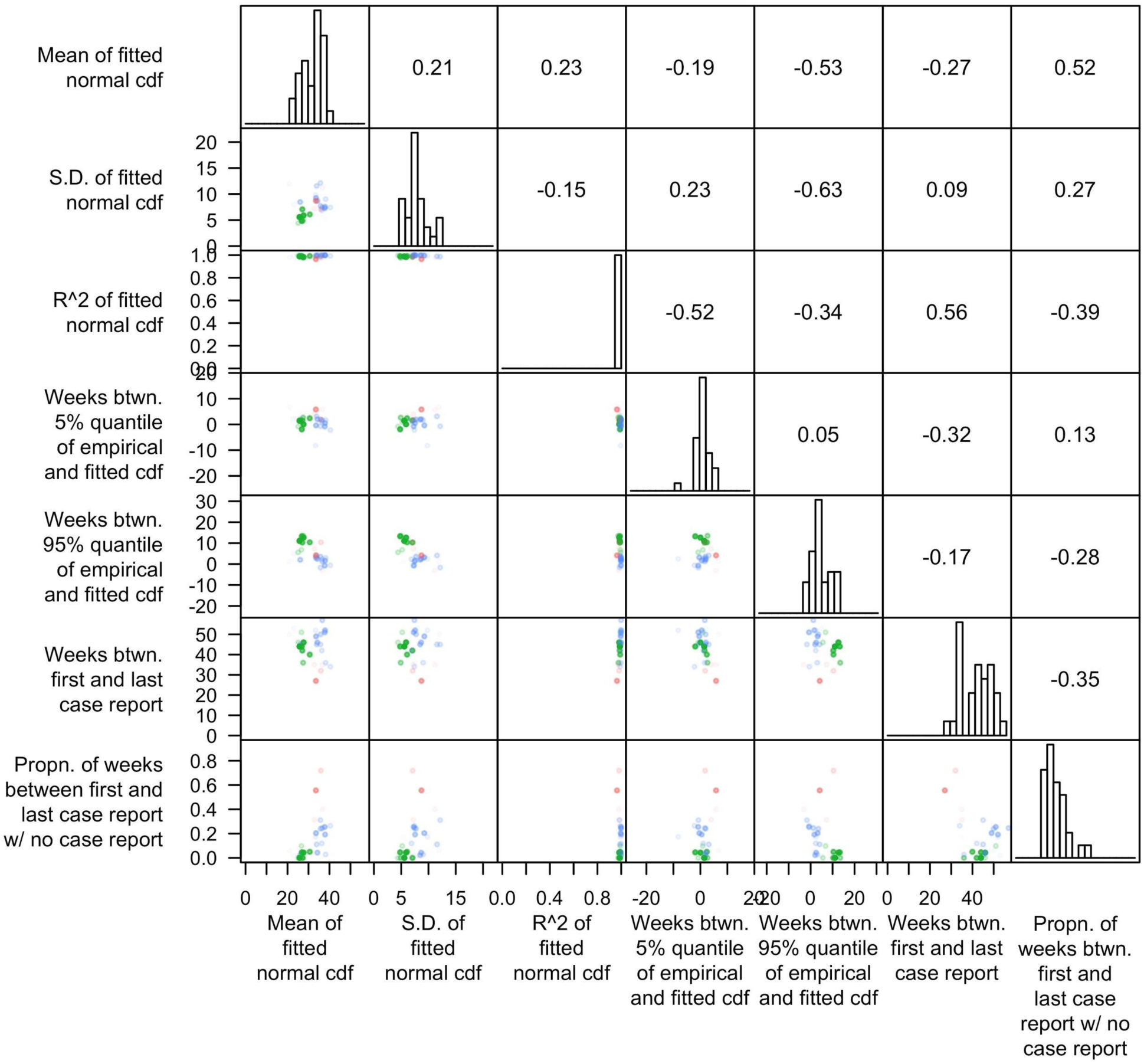
Pairwise plots of features of proportional cumulative incidence curves, with colors distinguishing group assignment of the departments into one of three groups. Histograms show the marginal distributions of the features, and numbers in the upper right half indicate pairwise correlation coefficients between each pair of features. The transparency of each point is inversely proportional to silhouette value.

**Figure S6.**
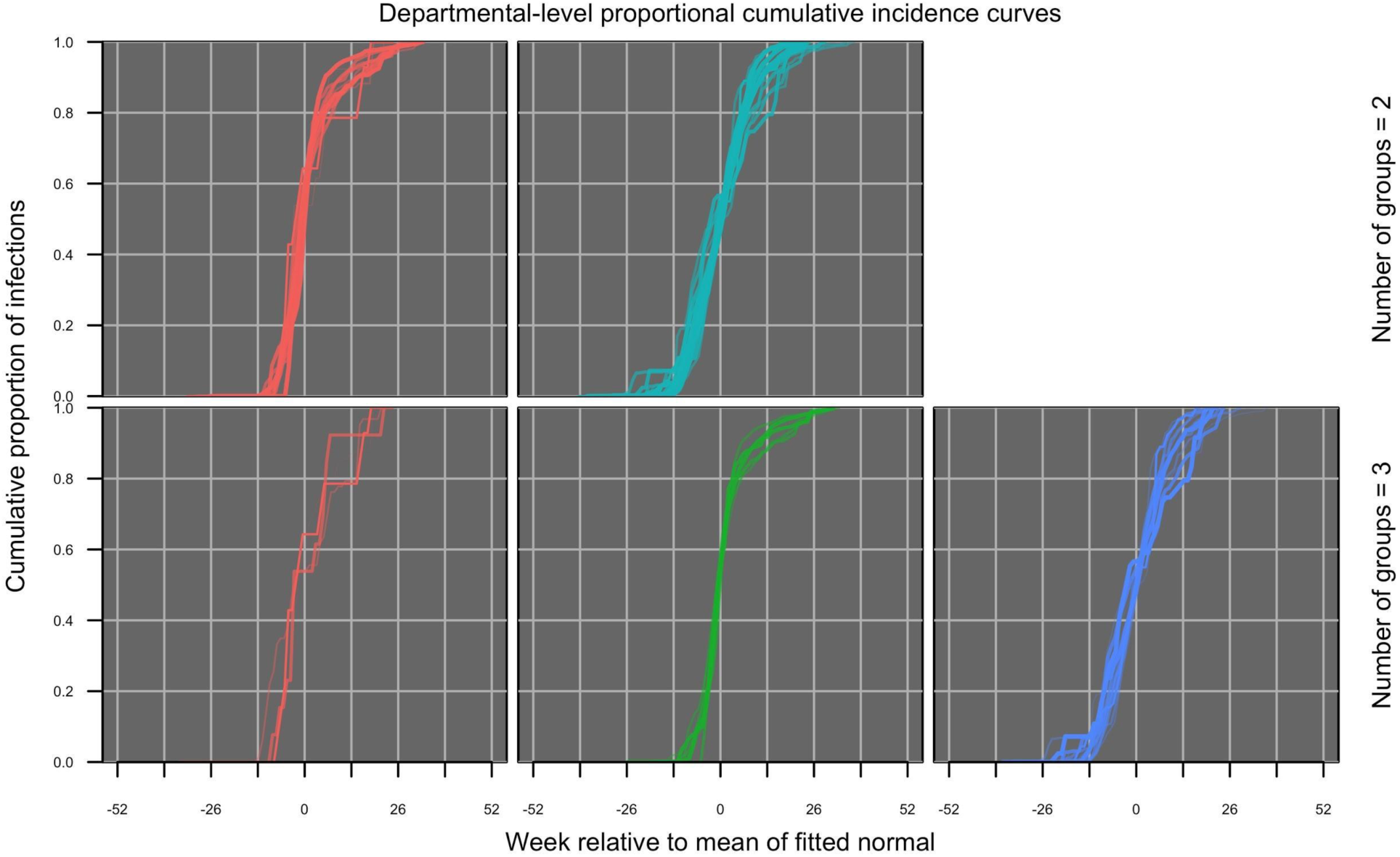
Proportional cumulative incidence curves at the departmental level with two (top) or three (bottom) groups. Within each row, groups are distinguished by color.

**Figure S7.**
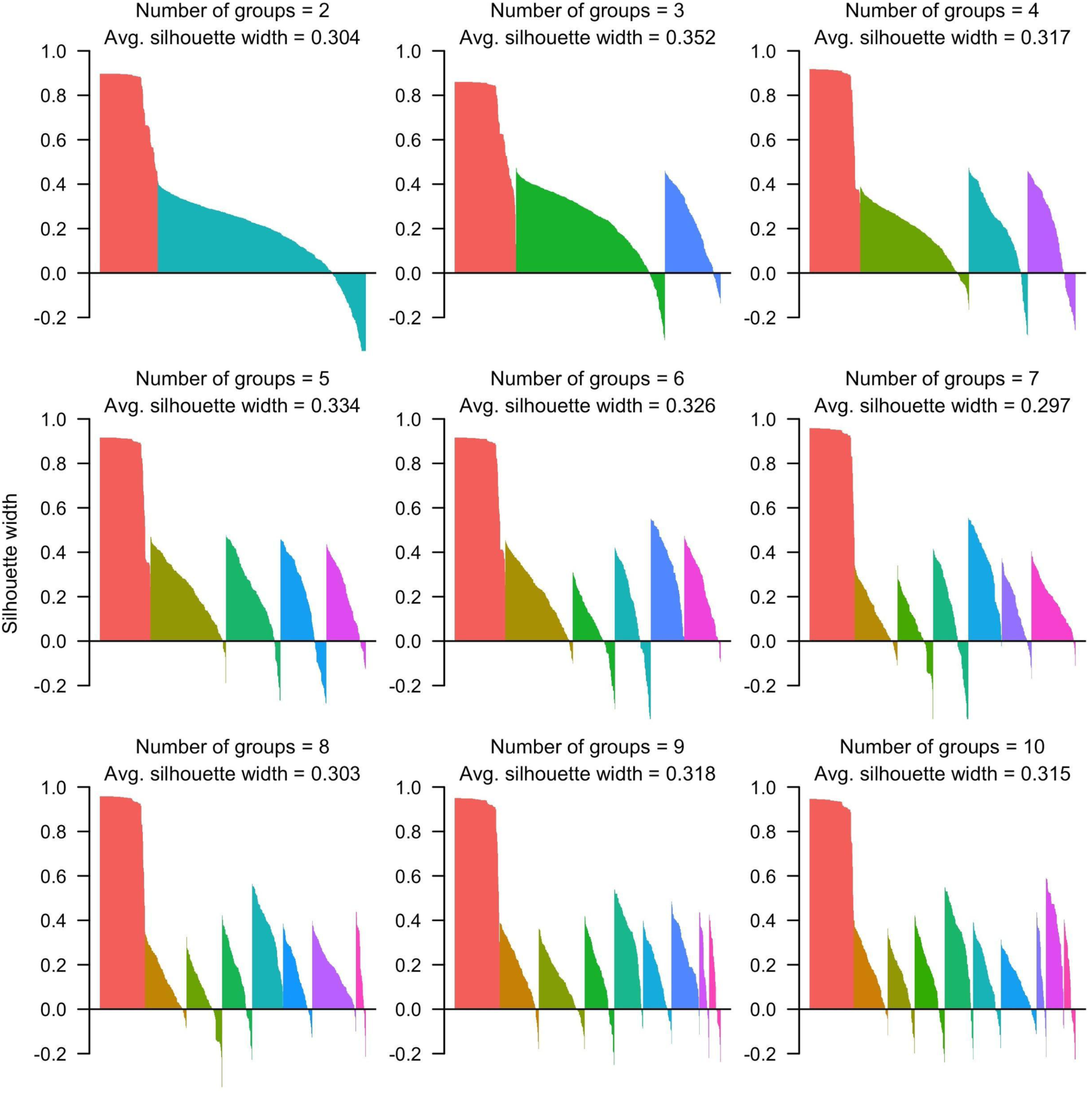
Silhouette plots at the municipal level for groups numbering two to ten obtained by partitioning around medoids. Each bar corresponds to the silhouette value of a given municipality according to the group assignments indicated by different colors in each panel. Higher average silhouette values indicate stronger clustering.

**Figure S8.**
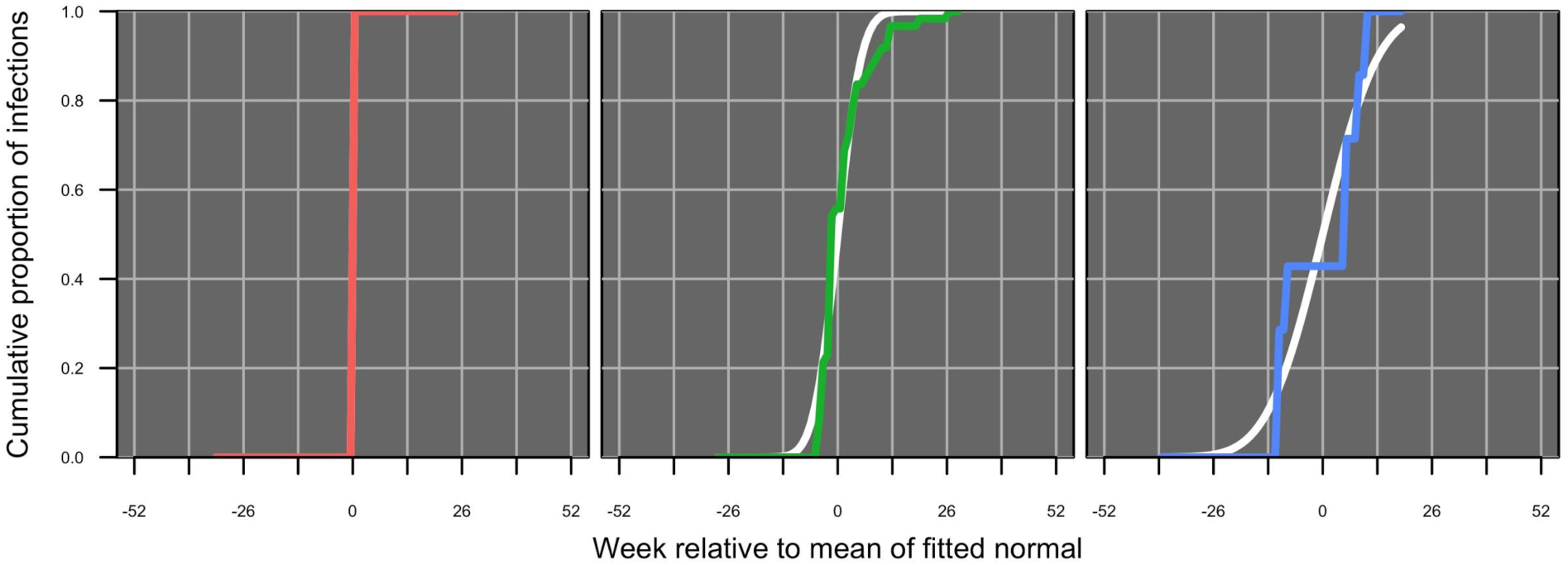
Proportional cumulative incidence curves (red, green, blue) shown against cumulative normal density curves (white) for medoids of the classification based on three groups at the municipal level.

**Figure S9.**
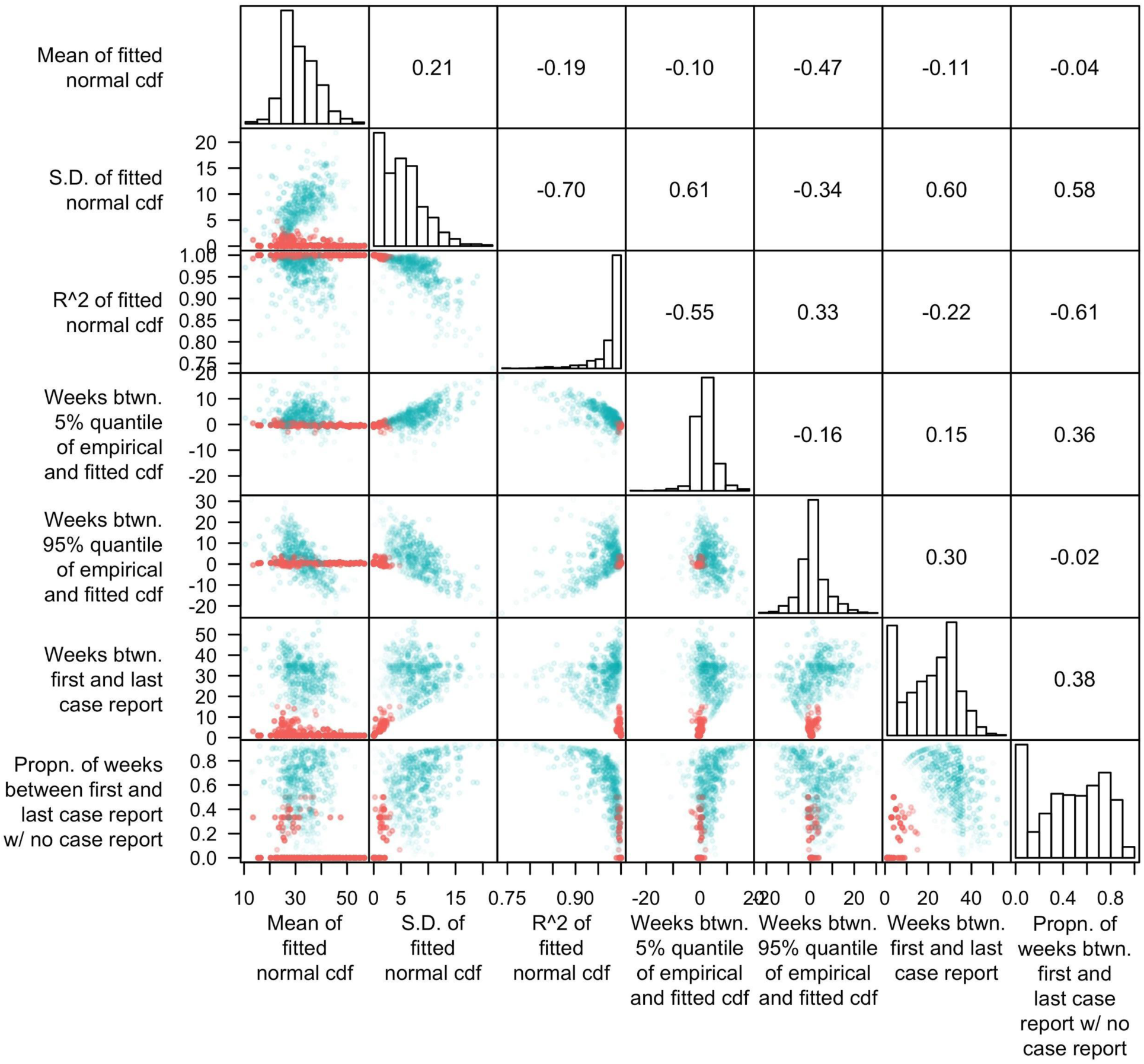
Pairwise plots of features of proportional cumulative incidence curves, with colors distinguishing group assignment of the municipalities into one of two groups. Histograms show the marginal distributions of the features, and numbers in the upper right half indicate pairwise correlation coefficients between each pair of features. The transparency of each point is inversely proportional to silhouette value.

**Figure S10.**
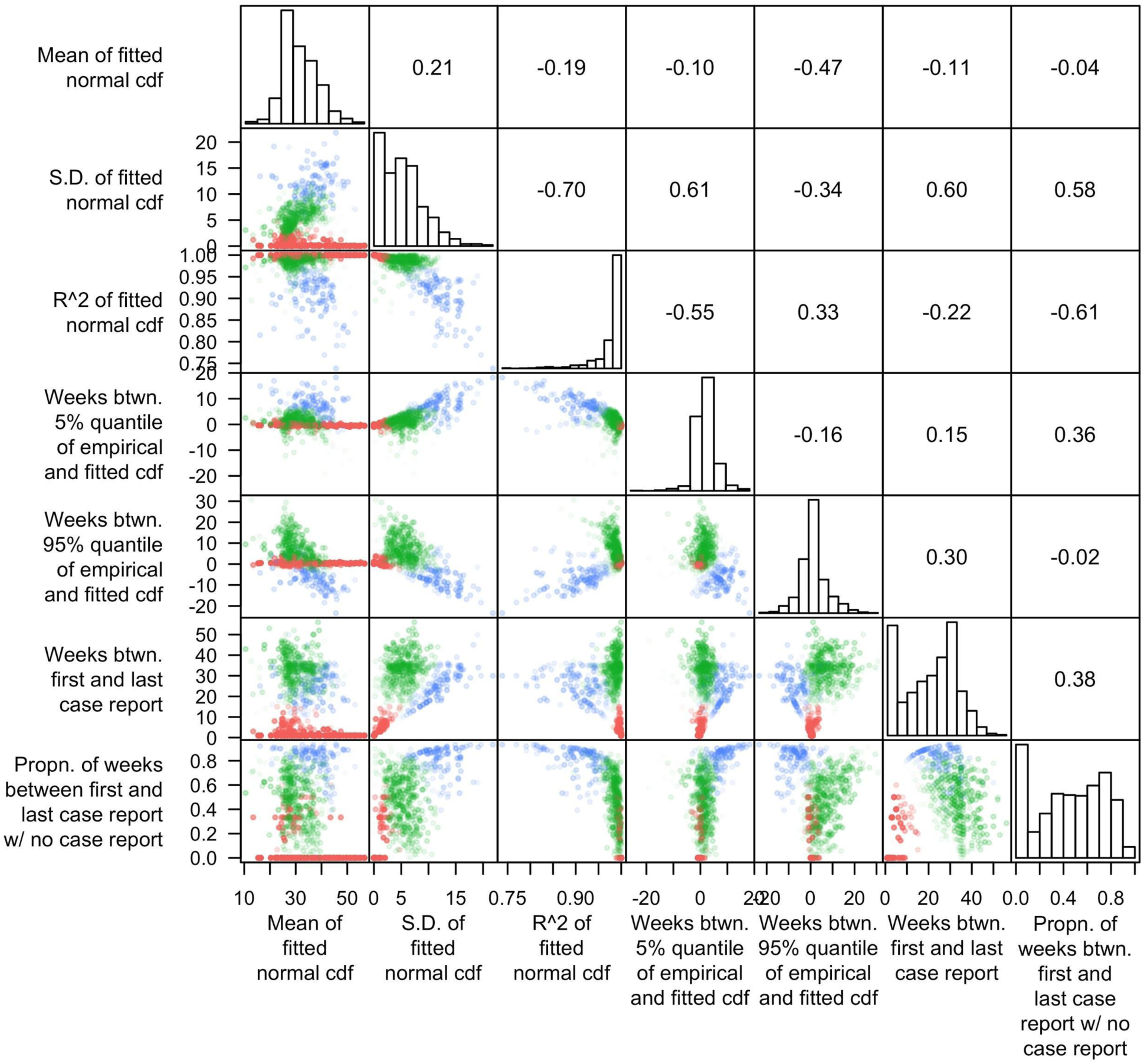
Pairwise plots of features of proportional cumulative incidence curves, with colors distinguishing group assignment of the municipalities into one of three groups. Histograms show the marginal distributions of the features, and numbers in the upper right half indicate pairwise correlation coefficients between each pair of features. The transparency of each point is inversely proportional to silhouette value.

**Figure S11.**
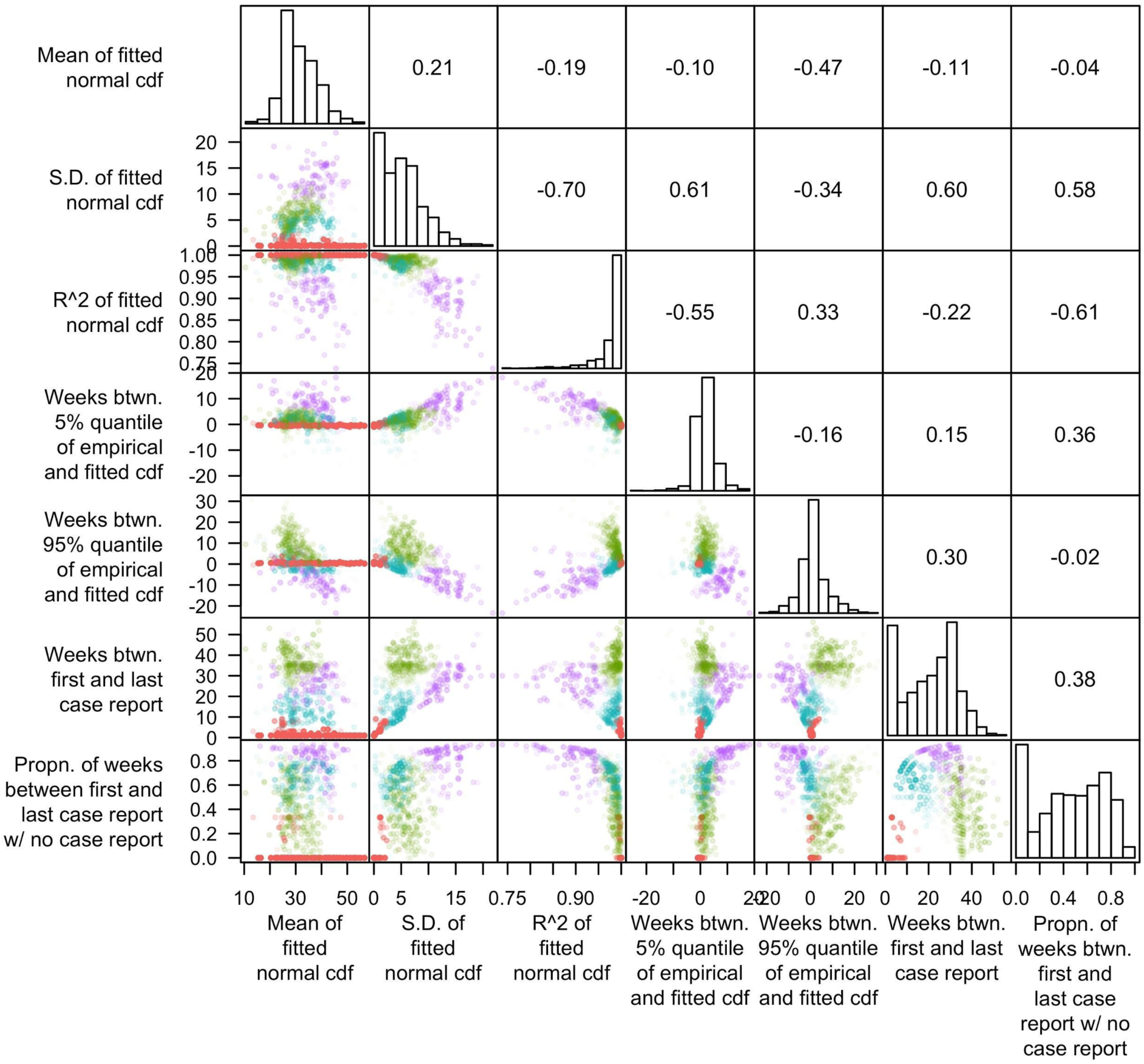
Pairwise plots of features of proportional cumulative incidence curves, with colors distinguishing group assignment of the municipalities into one of four groups. Histograms show the marginal distributions of the features, and numbers in the upper right half indicate pairwise correlation coefficients between each pair of features. The transparency of each point is inversely proportional to silhouette value.

**Figure S12.**
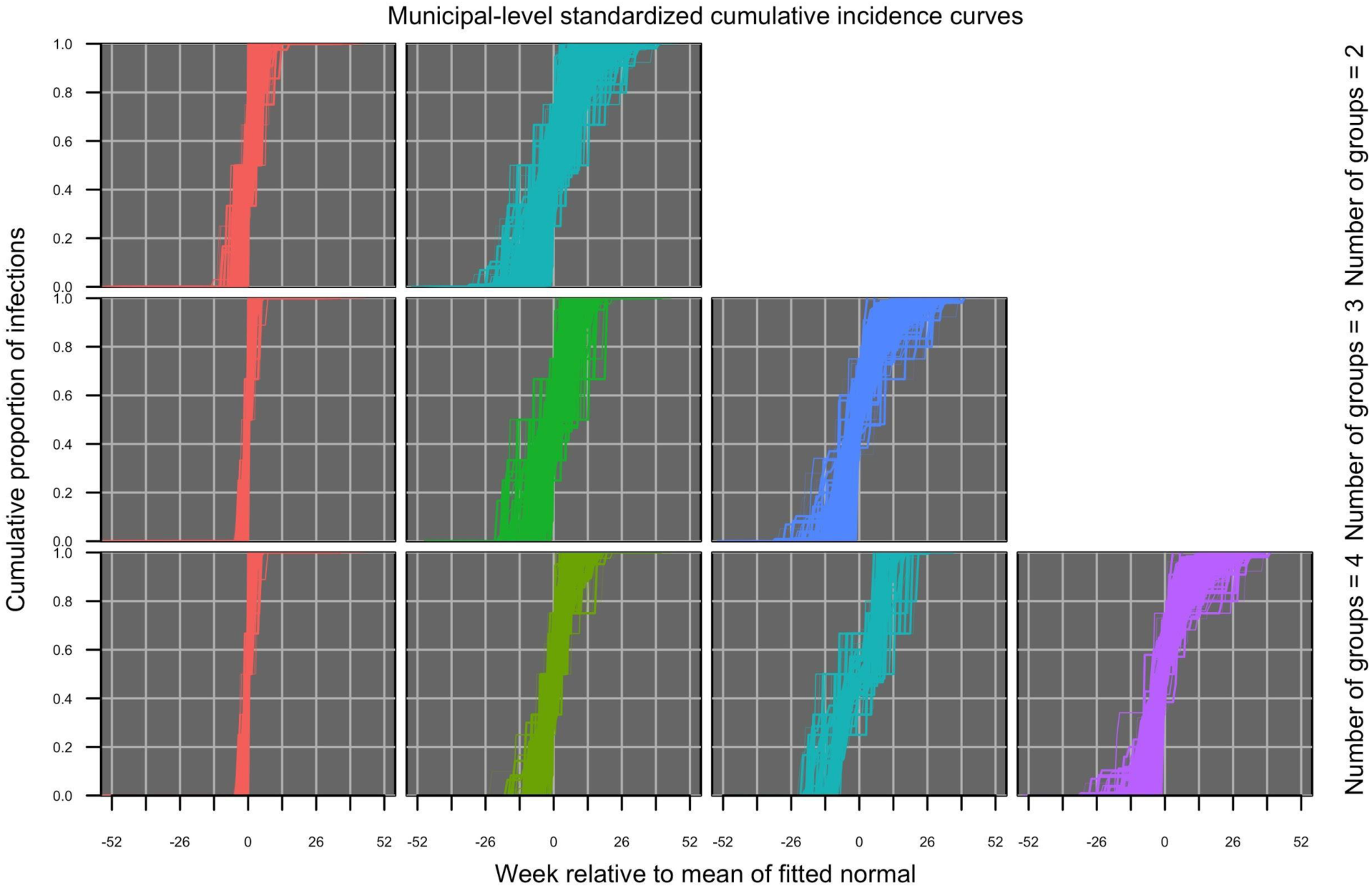
Proportional cumulative incidence curves at the municipal level with two (top), three (middle), or four (bottom) groups. Within each row, groups are distinguished by color.

**Figure S13.**
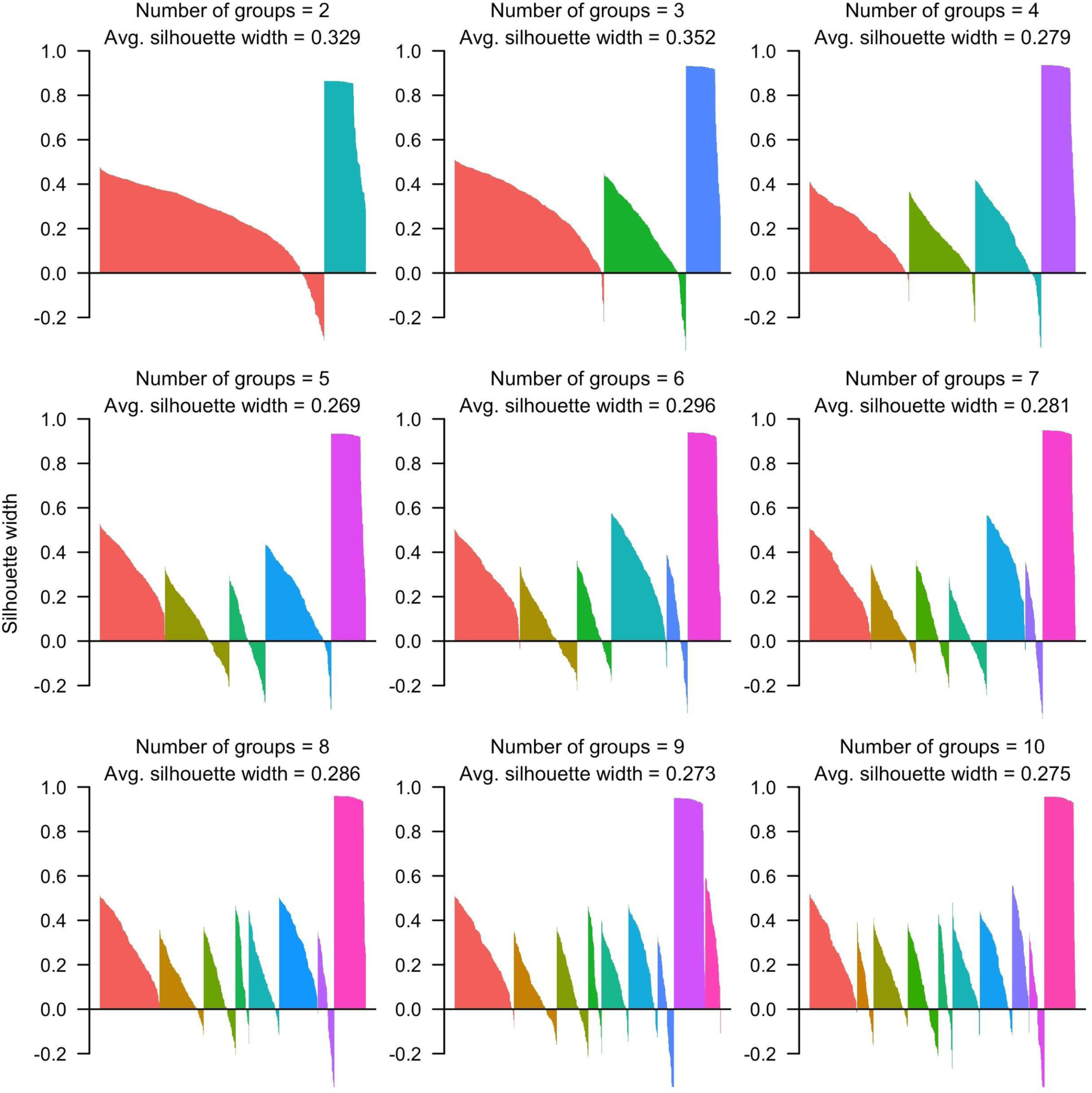
Silhouette plots at the municipal level based on a randomly selected simulated data set for groups numbering two to ten obtained by partitioning around medoids. Each bar corresponds to the silhouette value of a given municipality according to the group assignments indicated by different colors in each panel. Higher average silhouette values indicate stronger clustering.

**Figure S14.**
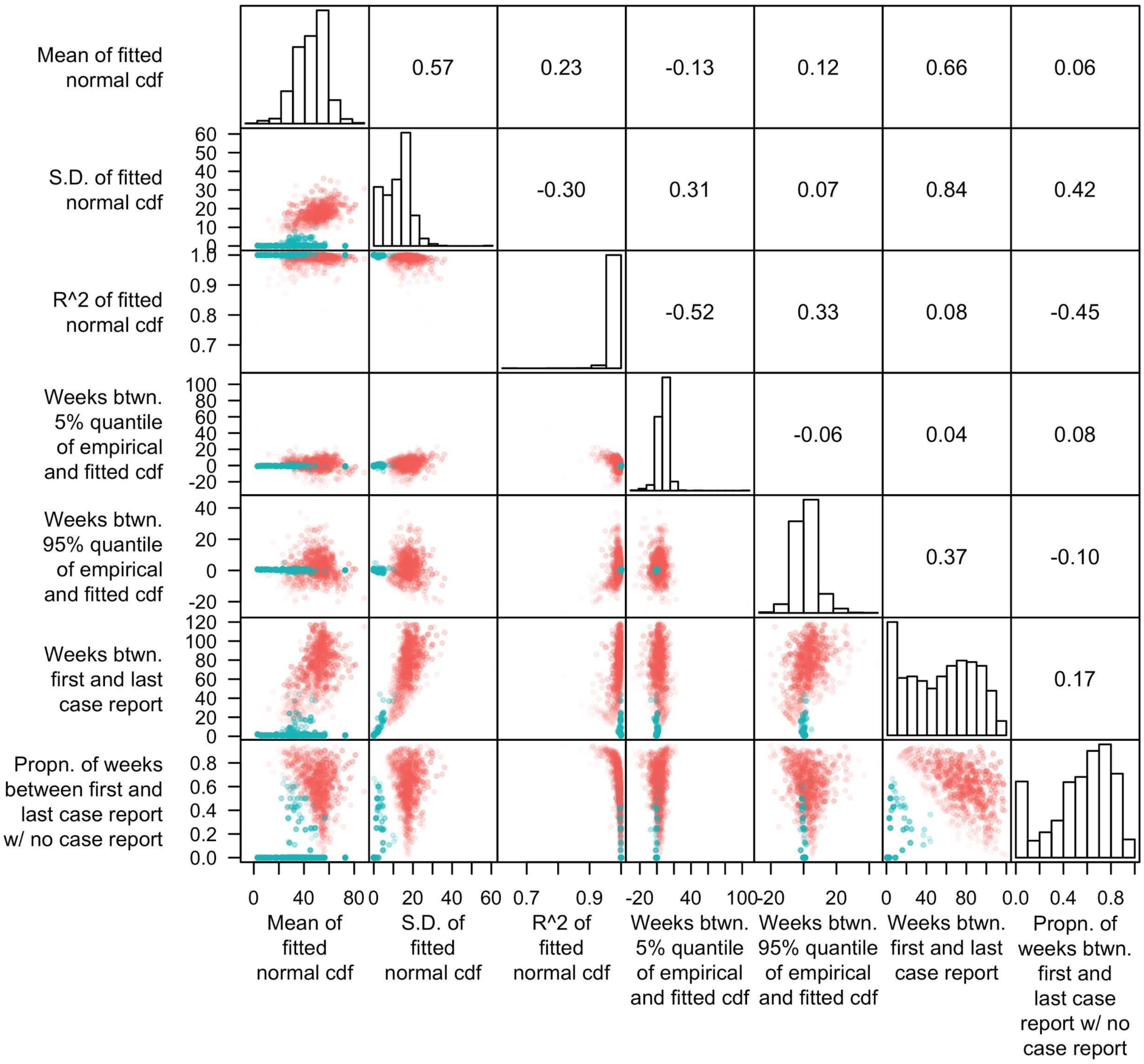
Pairwise plots of features of proportional cumulative incidence curves based on a randomly selected simulated data set, with colors distinguishing group assignment of the municipalities into one of two groups. Histograms show the marginal distributions of the features, and numbers in the upper right half indicate pairwise correlation coefficients between each pair of features. The transparency of each point is inversely proportional to silhouette value.

**Figure S15.**
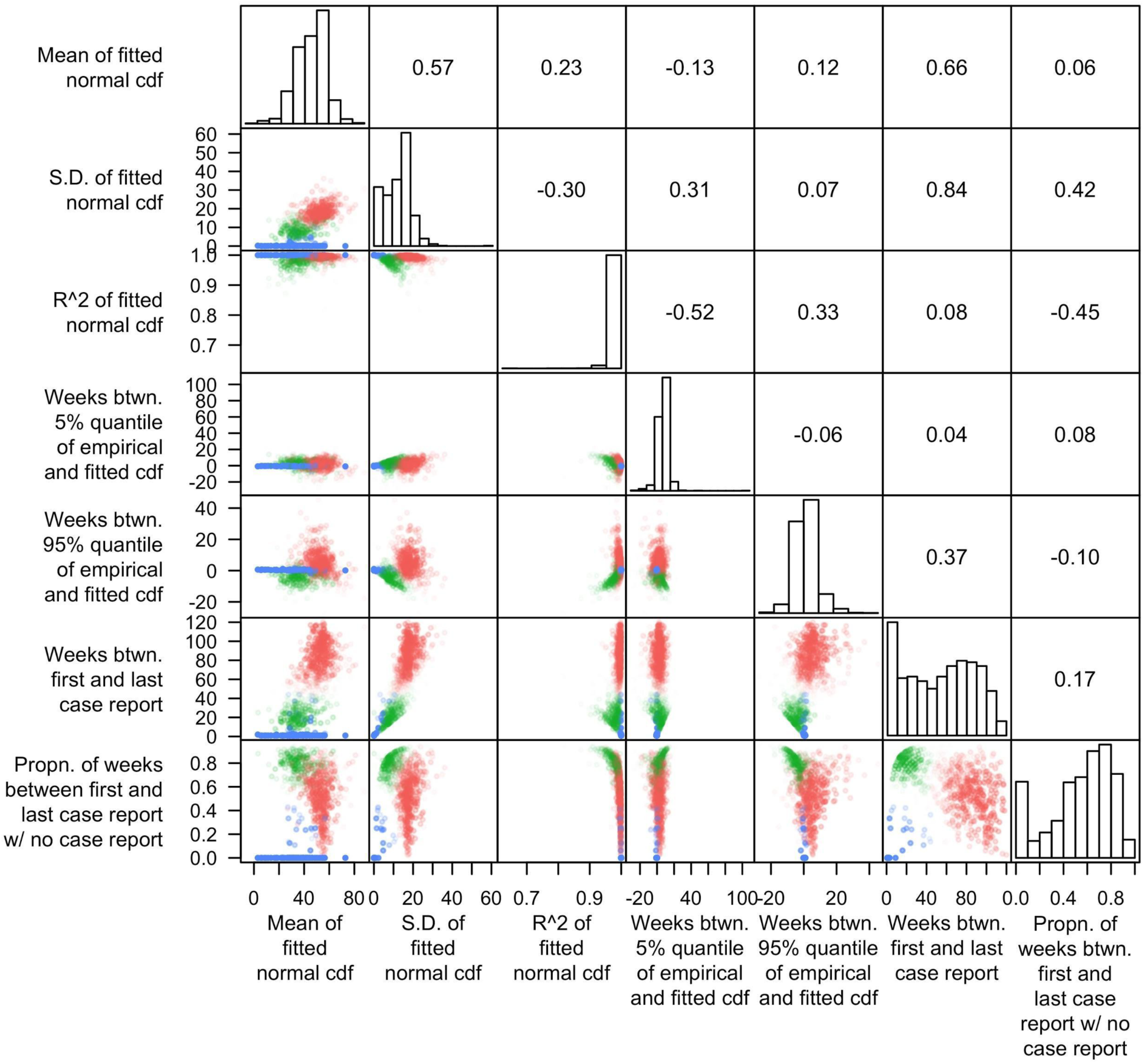
Pairwise plots of features of proportional cumulative incidence curves based on a randomly selected simulated data set, with colors distinguishing group assignment of the municipalities into one of three groups. Histograms show the marginal distributions of the features, and numbers in the upper right half indicate pairwise correlation coefficients between each pair of features. The transparency of each point is inversely proportional to silhouette value.

